# Widespread Recruitment of ParA-like NTPases into Prokaryotic Immunity reveals a diversity of anti-selfish element strategies

**DOI:** 10.64898/2026.10.19.760177

**Authors:** Huan Li, A. Maxwell Burroughs, L Aravind

## Abstract

The SIMIBI subgroup within the GTPase class of P-loop NTPases, including ParA/MinD, is best known for chromosome segregation, cell division, and macromolecular localization. Its role in immunity remains largely unexplored relative to TRAFAC-subgroup GTPases and other NTPases. Using systematic sequence, structure, and comparative genomics analysis, we surveyed the ParA clade for roles in immunity. ParA underwent at least two major, independent transitions from core cellular functions to prokaryotic immunity, producing two large "conflict" branches with diverse, uncharacterized systems, plus smaller, restricted immune lineages arising sporadically outside these branches. Across 28 conflict systems, ParA couples to a wide range of partners, including nuclease, NADase, and peptidase effectors, novel sensor domains such as a variant HeH domain, and apoptosis-inducing modules reminiscent of eukaryotic programmed cell death; several systems are enriched in multicellular bacteria, and a subset was transferred to eukaryotes on multiple occasions. Newly characterized ParA partner domains, including a nucleotide cyclase, a TIM-barrel domain, START, and permuted FHA domains, further define new anti-selfish-element systems independent of ParA. Together, these findings establish ParA as a recurrently co-opted "fulcrum" NTPase in prokaryotic immunity, predicted to act as an interceptor of invasive nucleic acids, a multimerizing effector, and a switch regulating other immune effectors.

## Introduction

Nucleotides are foundational molecules of life, functioning as building blocks of genetic material, energy carriers, first and second messengers in signaling, and substrates and products in most key biochemical pathways. The capacity to catalyze the key cellular energetic reaction, namely the hydrolysis of nucleotide triphosphates (NTPs), has evolved in several protein domain superfamilies with distinct folds [1]. The most widely distributed and abundant of these is the P-loop NTPase fold [2–7]. The P-loop NTPase fold is characterized by a three-layered α/β sandwich, featuring a mostly parallel core β-sheet composed of five universally conserved strands (β1 to β5) on either face of which are packed α-helices that are interleaved with the strands [6]. At the sequence level, P-loop NTPases possess two characteristic signatures, the Walker A (or the P-loop) and Walker B motifs. The Walker A motif, which binds the triphosphate moiety of NTPs, typically appears as GxxxxGK[ST] and is located in the loop between strand β1 and helix α1 [8]. The Walker B motif, characterized by the hhhh[D] signature (where h is a hydrophobic residue and the aspartate contributes to Mg²⁺-binding), encompasses strand β3 [9]. P-loop NTPases harness the free energy of NTP hydrolysis to participate in an astonishing array of biochemical mechanisms by acting as (i) macromolecular switches, or (ii) motors that rearrange contacts between macromolecules, or (iii) phosphotransferases that transfer phosphates or other moieties from NTPs to target molecules [8, 10].

GTPases, a major monophyletic assemblage within the P-loop NTPase superfamily, in turn group with the P-loop Kinases to form the (Kinase-GTPase) KG division of the superfamily [3]. This division is characterized by a distinctive structural arrangement in which the β1 strand associated with the Walker A motif directly hydrogen bonds with the β3 strand bearing the Walker B motif, yielding a 54132 β-sheet topology (Figure 1A) [3]. Further, in the GTPase assemblage, the β5 strand is ancestrally followed by the [N/T]KxD motif [11], which is central to GTP selectivity. However, this motif has degenerated on multiple occasions across the GTPase assemblage, resulting in clades that retain the ancestral GTPase fold but have transitioned to ATPase activity. Examples of these include the myosin-kinesin, ParA, and BioD families [3]. All GTPases belong to one of two higher-order subgroups. The first subgroup, termed TRAFAC after its founding members, the translation factors, features the transformation of the ancestral helix between the strands β2 and β3 into an additional antiparallel strand (βB) inserted into the core sheet [3]. In contrast, the second subgroup, termed SIMIBI, has an additional α-β unit inserted after strand β2, thereby augmenting the sheet with an additional parallel strand (βB; Figure 1A). However, in both subgroups, this region bears a conserved "sensor" residue (e.g., DxD for ParA, Figure 1A) that discriminates between the NTP-and NDP-bound states.

**Figure 1.**
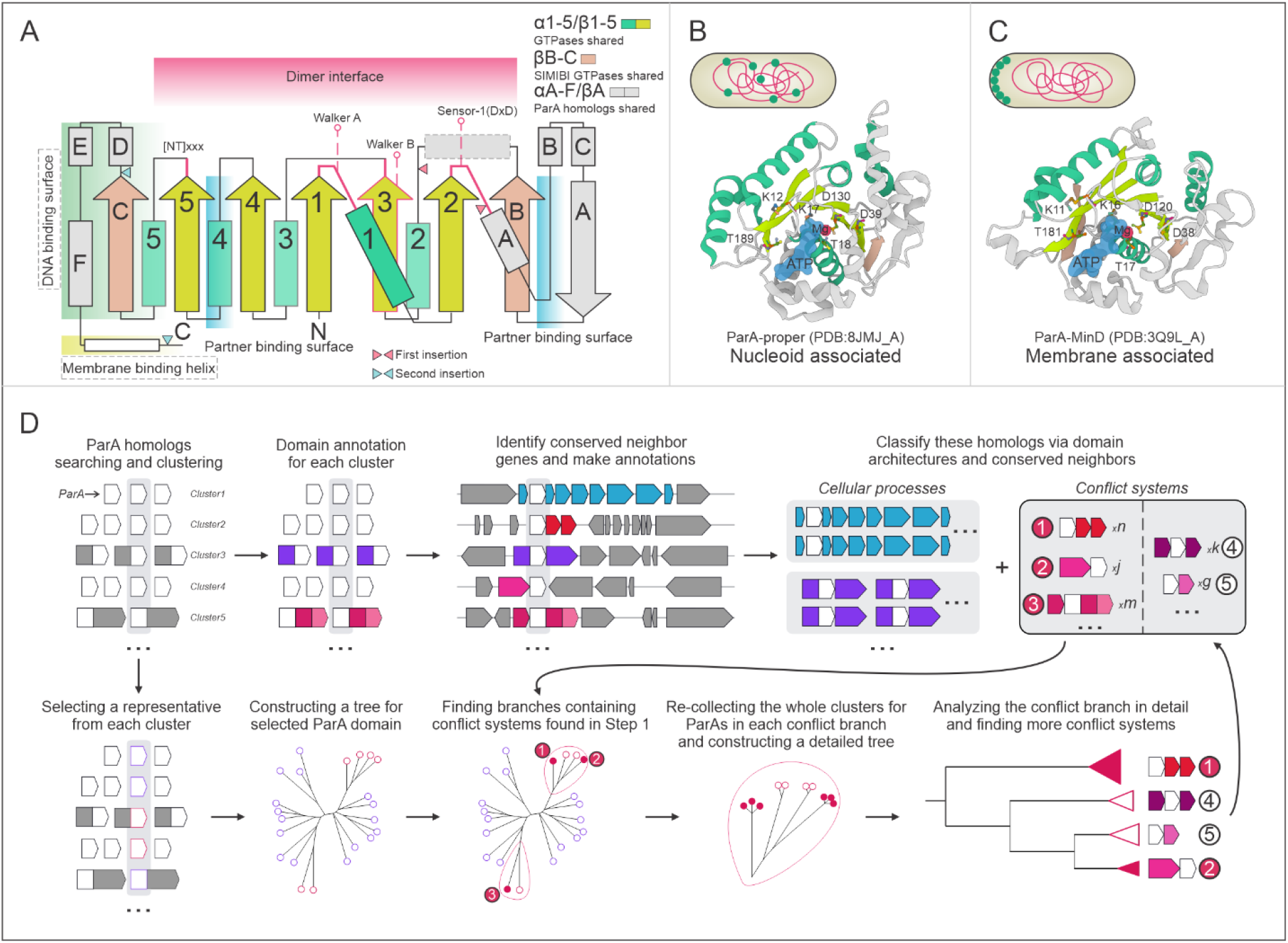
(A) Topology of the ParA family, including structural elements, conserved motifs, and binding surfaces. (B) Cartoon representation of the ParA domain in the classical ParABS system, which has been shown to bind the nucleoid. The PDB ID, conserved residues, Mg²⁺ ion, ATP substrate, and a schematic illustrating the subcellular location of the ParA domain are shown. (C) Cartoon representation of the MinD-type ParA domain in the MinCDE system, which is localized at the cell membrane. (D) Workflow for identifying novel biological conflict systems based on the ParA domain in this study. **Alt text:** Figure with subfigures labelled A to D, showing the topology and cartoon structures of the ParA protein family in the ParABS and MinCDE systems, and a workflow for identifying biological conflict systems.

GTPases participate as molecular switches in a wide range of fundamental cellular processes, including but not limited to (i) translation, such as initiation and elongation factors [12]; (ii) membrane localization and secretion (e.g., the signal recognition particle GTPases) [13]; (iii) membrane trafficking (e.g., dynamins) [14, 15]; (iv) signal transduction (e.g., Ras) [16]; (v) archaeo-eukaryotic (e.g., septins) [17] and bacterial (e.g., ParA/MinD) cell division [18]; and (vi) metabolism (e.g., BioD [19] and NifH [20]). In recent years, an explosion of interest in immune processes in both prokaryotes and eukaryotes [21–27] has been accompanied by a growing body of new insights into the distinct families of P-loop NTPases in these processes, including GTPases, ABC ATPases [6], STAND NTPases [28–31], and helicases [32, 33]. TRAFAC subgroup GTPases have been studied as regulatory switches and drivers of membrane reorganization in numerous immune-related pathways, most notably, the antiviral representatives of the AP-GTPase, GIMAP, Septin, Mx/Dynamin-like, and the recently characterized bacterial DO-GTPase clades [28, 31, 34]. However, to date, the roles of members of the SIMIBI subgroup in these processes remain less understood.

The ParA and closely related SIMIBI GTPase class lineages [3] have previously been best characterized in ensuring the correct positioning of DNA or proteins within cells to mediate chromosome segregation and correct macromolecular localization [18]. Previously characterized roles of ParA clade NTPases include: (i) ParA proper in the ParABS system regulating chromosome segregation and plasmid partition [35]; (ii) MinD in preventing the formation of the divisome, the apparatus controlling bacterial cell division, at the poles of bacterial cells [36]; (iii) FlhG in regulating the correct location and number of flagella [37]; (iv) ParC in recruiting components of chemotaxis clusters to newly forming cell poles [38]; (v) VirC1 in directing the spatial localization of the transferred DNA strand to the cell pole during conjugation [39]; (vi) McdA in facilitating the spatial organization of carboxysomes, a specialized bacterial microcompartment that enhances the efficiency of CO₂ fixation [40]. In recent studies, we observed members of the ParA clade of the SIMIBI subgroup to be involved in predicted prokaryotic immune systems with potential connections to apoptotic mechanisms [28]. The presence of ParA clade domains in a predicted bacterial immune system raised the possibility that unrecognized ParA homologs might be involved in uncharacterized biological conflict systems, prompting us to systematically explore these SIMIBI NTPases for such roles.

In this study, we comprehensively collected ParA clade homologs involved in biological conflicts broadly defined by immune processes using a systematic sequence, structure and comparative genomics protocol. Our investigation revealed that within the ParA clade, there have been two major evolutionary transitions from roles in fundamental cellular processes to roles in immunity, giving rise to two ’conflict’ branches that contain diverse, previously uncharacterized systems. In addition, we also established the repeated sporadic emergence of phyletically more restricted ParA-based immune systems that are not included in these two branches. Finally, we also discovered a diverse array of additional prokaryotic immune systems by analyzing novel protein domains contextually associated with ParA clade proteins in predicted immune systems. These investigations unveil a world of NTPases from the SIMIBI subgroup involved in potential immune processes and predict novel molecular mechanisms in host-virus biological conflicts.

## Materials and Methods

### Sequence analysis

Comprehensive sequence searches were performed using the PSI-BLAST program [41], with an e-value cut-off of 0.001, against the NCBI non-redundant (nr) protein database clustered down to 50% sequence similarity via the MMseqs2 program [42]. The identified homologs were further clustered by the BLASTCLUST program (https://ftp.ncbi.nih.gov/blast/documents/blastclust.html) to select representative sequences for the Multiple Sequence Alignment (MSA), which was generated using FAMSA [43] and MAFFT programs [44] followed by the computation of a consensus for each family from the MSA [45]. The MSA profiles and consensus patterns for the ParA domain and other domains identified in this study are provided in the Supplementary Data. The HMMER package [46] was used to build HMM profiles for the novel domains based on the MSA, which were then used for downstream contextual analyses (see below). HMM sequence logos were generated via the logomaker program [47]. Sequence network clustering was performed using the CLANS program [48].

### Contextual analysis

Gene neighborhood analysis [49] was used to identify novel systems based on the ParA domain or other domains, by filtering for their conserved neighboring genes. Specifically, homologs of the target domain identified by PSI-BLAST were used to retrieve their upstream and downstream genes (five genes in each direction) from the NCBI RefSeq genomes database. All proteins in the neighborhood were annotated for domain architectures using HMMSCAN [46] and RPS-BLAST against the Pfam database [50] and the above-mentioned collection of new profiles. The neighborhood proteins were then clustered based on sequence similarity using the BLASTCLUST program. Clusters of conserved neighbors were defined by two criteria: (1) they are within 50 nucleotides of the target gene in genomic loci; and (2) the associations are conserved across at least two different bacterial phyla. Contextual networks based on domain architectures and neighborhoods were generated using the Gephi software [51].

In addition, we further incorporated a phylogenetic inference, which is more accurate than clustering based solely on pairwise sequence similarity [52], as a downstream step to identify rarer contextual linkages. Specifically, homologs of the target protein were clustered using BLASTCLUST, and one representative sequence was selected from each cluster. The representative sequences were then used to build an overall phylogenetic tree for the target domain, and novel biological conflict systems previously identified via gene neighborhood analysis were mapped onto the tree. Branches containing novel biological conflict systems were then extracted, and their representative sequences were used to trace back the clusters generated by BLASTCLUST to which they belong. The clusters were then used to build a detailed phylogenetic tree for that conflict-containing branch, which was used to identify additional novel biological conflict systems. This additional step is especially effective for investigating domains like ParA, given that it is a highly abundant family in the database (>50,000 homologs in NR50) and encompasses both fundamental cellular processes and biological conflict systems.

### Structural analysis

Experimentally determined protein structures were obtained from the PDB database [53]. Structural models presented in this study were predicted using the AlphaFold server [54]. The structural models and their associated PAE matrices were used to determine domain boundaries, and only domain structures with overall reliable pLDDT scores [55] and pTM score > 0.5 [56] were selected for presentation and inference. Structure-structure similarity searches with DALI [57] and Foldseek [58] were used to identify distantly related homologs [59–76]. Renderings of the protein structures were generated using Mol* viewer [77].

### Phylogenetic analysis and multicellular enrichment test

Representative sequences of the ParA domain from each system, including those involved in basic cellular processes and immune systems, were used to build the phylogenetic tree using the IQ-TREE software [78]. The phylogenetic tree was inferred using the LG+C60+F+G+PMSF substitution model [79], with branch support evaluated using 1,000 ultrafast bootstrap replicates and 1,000 SH-aLRT replicates. The phylogenetic tree file and the ParA MSA used to construct the tree are provided in the Supplementary Data. Visualizations of the phylogenetic tree were generated using the TreeViewer software [80].

The taxonomic information for ParA homologs was extracted from NCBI GenPept files. A database of 8052 representative prokaryotes was compiled from NCBI RefSeq and used to analyze the phyletic distribution patterns of ParA-based systems. These organisms were scored for their habit (multicellular, unicellular and unknown) and used to calculate the enrichment of each system in multicellular organisms. The hypergeometric distribution test was used to test significance, as described earlier [28], with species in the database set as the background frequency and labeled as TRUE, FALSE, and NA for the multicellularity flag.

## Results and discussion

### Defining the conserved structure and sequence core of the ParA clade

We first examined known members of the ParA clade using multiple sequence alignments to identify their characteristic sequence features [3]. These include (Figure 1A-C): (i) a distinct Walker A motif (KGGxGK[ST]), which features an additional conserved lysine; (ii) a Walker B motif (DhhhhD), containing an additional conserved aspartate at the beginning (this feature is shared with other members of the SIMIBI subgroup); (iii) a degenerate [NT]KxD motif retaining only the first N, consistent with ParA having lost its GTP preference and binding ATP as a substrate [18]; (iv) a DxD motif in the sensor region at the end of the core strand β2, which additionally contributes to Mg²⁺ binding along with the Walker B motif [81, 82].

We then examined experimentally determined and AF3-generated structures to identify ParA-clade-specific structural elements beyond those shared by most SIMIBI GTPases. These include helical insertions or terminal extensions (helices αA to αF), and an antiparallel β-strand (βA stacking alongside βB (Figure 1A)). Next, we surveyed prior studies to map the known interaction surfaces of the ParA SIMIBI NTPase domain relative to its defining structural elements [18]. These include: (i) the surface mediating ParA dimerization, which is formed by the edge of the core sheet marked by the C-termini of its constituent β-strands. This surface incorporates the first conserved lysine of the ParA Walker A motif [81, 82] (Figure 1A). (ii) The surface defined by the C-terminus of the domain, which mediates binding to specific macromolecular structures associated with localization. Characterized members of the ParA family can be broadly categorized into two types based on their macromolecular associations in this region (Figure 1BC) [18]. The first category, including ParA proper [81], McdA [83] and ParC [84], interacts with nucleoid DNA through several basic residues located around core helix α5 and additional helices αD-αF. The second category, including MinD [82], FlhG [37], and VirC [39], associates with the cell membrane via a further amphipathic helix at the extreme C-terminus [85]. (iii) The surface formed in a ParA dimer by the insert in the core sheet between β2 and β3 in one monomer and the core helix 4 in the other monomer [81, 82, 86] mediates interaction with a partner protein, such as ParB in the ParABS system and MinE in the MinCDE system. These partner proteins facilitate the disassociation of their cognate ParA from their macromolecular associations, a necessary step for the dynamics of ParA-based systems.

### Computational pipeline for the identification of ParA clade proteins involved in immune functions

We first comprehensively collected homologous sequences using known ParA domains as PSI-BLAST queries and separated them from other SIMIBI domains using the above-described features (Figure 1D, upper panel; see Methods). After annotating these homologs and their conserved gene-neighbors using RPS-BLAST and HMM searches, we grouped them into distinct systems based on their domain architectures and gene-neighborhood partners. To separate ParA homologs with roles in immunity from the ‘institutionalized’ versions participating in conserved cellular processes, we leveraged the characteristic hallmarks of biological conflict systems [49] and contextual information from conserved gene neighborhoods [87, 88] (Figure 1D, top panel illustrates the basic protocol). This approach is particularly effective for identifying systems whose components are abundant in the sequence database.

Beyond this, we incorporated a second step based on phylogenetic clustering (the bottom panel of Figure 1D shows this step). ParA domains collected via PSI-BLAST were clustered based on sequence similarity to reduce redundancy from closely related sequences (see methods), and the non-redundant set was used to construct a phylogenetic tree. The potential immune versions identified via gene neighborhood analysis were then mapped onto the tree, and the clades containing them were isolated. For each such clade, clade-specific trees were constructed using all sequences from the original clusters (Figure 1D, upper-right panel). These were then analyzed in detail based on their contextual connections to identify additional biological conflict systems. This step was effective at identifying systems that are sparsely represented in the database, particularly those with unexpected or atypical domain associations. The compiled taxonomic distribution data of these systems are reported in Supplementary Fig. S1. This data was also used to test each system for statistical overrepresentation in bacterial taxa with particular habits or morphological features (e.g., multicellularity; see below for details [89, 90]).

### Diversification of ParA NTPases reveals two major independent transitions from core cellular processes to immune functions

The tree of the ParA NTPases revealed two primary clades, both found across diverse prokaryotic lineages, and associated with distinct conserved gene neighborhoods (Figure 2A, B). The first clade, PriB1 (Primary Branch 1), includes ParA homologs from ParABS bacterial chromosome segregation systems and versions mediating the localization of chemotaxis receptors and carboxysomes. The second clade, PriB2, includes members associated with flagellation patterns, conjugation, and bacterial divisome organization. The distinctiveness of PriB1 and PriB2 is also supported by the Hidden Markov model logos of their conserved motifs (Figure 2C). Notably, in the Walker A region, PriB1 contains a conserved asparagine at the N-terminal-most position, whereas PriB2 contains a conserved serine at the equivalent position. Similarly, in the sensor-1 motif, PriB1 is marked by a highly conserved glutamine two positions downstream of the characteristic DxD, which is absent in the PriB2 clade (Figure 2C).

**Figure 2.**
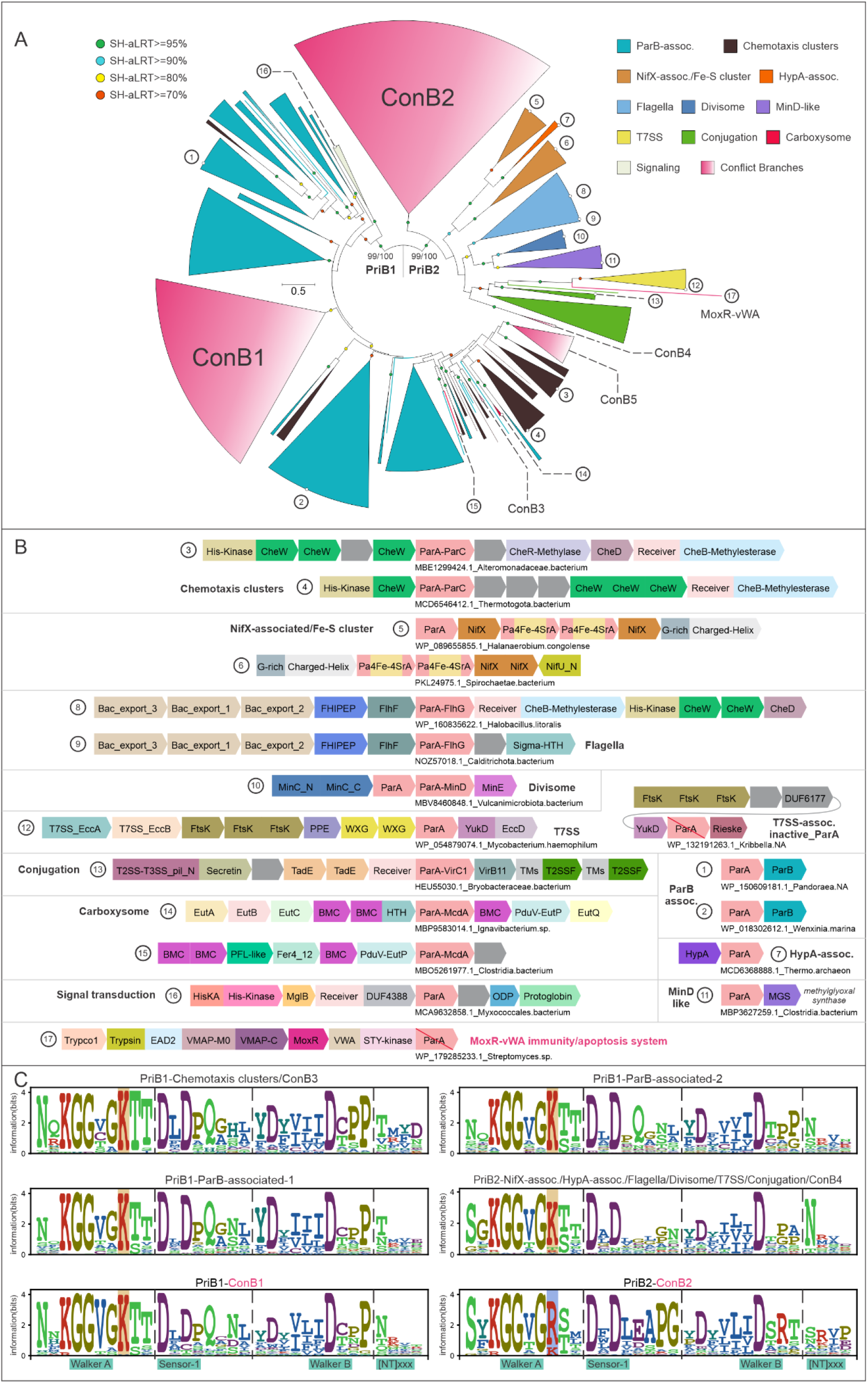
(A) Phylogenetic tree of the ParA family, generated using representative ParA homologs involved in basic cellular processes and immune systems identified in this study. Clades are colored according to their associated systems, with the immunity-associated branches (ConB1-ConB5) highlighted in red. SH-aLRT support values ≥ 70 are indicated by circles on the branches and color-coded by value. For the primary branches (PriB1 and PriB2), both ultrafast bootstrap and SH-aLRT support values are shown. (B) Representative depiction of ParA-associated basic cellular systems and the previously described MoxR-vWA immunity/apoptosis system [28]. Each arrow represents a gene within the genomic locus, and the encoded domains are shown with distinct colors. Genes, domains, and intergenic distances are not to scale. The NCBI accession ID of the ParA protein and the corresponding organism name are provided for each example. The number assigned to each panel corresponds to that in 2A. (C) HMM sequence logos of conserved motifs (Walker A, Sensor-1, Walker B, and [NT]xxx) across six phylogenetically distinct groups of ParA domains. The second lysine residue of Walker A and its arginine substitution in ConB2 are highlighted. **Alt text**: ParA family systems and conserved features. Figure with subfigures labelled A to C, showing a phylogenetic tree of the ParA family with immunity-associated clades highlighted, genomic locus diagrams of ParA-associated cellular and immune systems, and sequence logos of conserved motifs across six ParA groups.

Although not the focus of this study, a closer examination of the two clades furnished several novel insights into the evolution of systems involved in “housekeeping” cellular processes. First, our gene-neighborhood analysis, coupled with the tree, suggests that recruitment of ParA domains participating in the organization of chemotaxis receptors likely occurred multiple times independently within the PriB1 clade (Figure 2A, B, panels 3 and 4). PriB2-clade ParA homologs involved in flagellar patterning, i.e., FlhG, also appear to be incorporated in some cases into genomic loci with chemotaxis signaling genes (Figure 2B), consistent with the functional coupling between chemotaxis signaling and flagellar patterning [91]. We also observed that in certain cases, PriB1 clade ParA domains displace the Ras-like GTPase MglA in signaling systems centered on the MglA-MglB switch that regulates gliding motility in *Myxococcus* and related bacteria [92, 93] (Figure 2B). This indicates that the ParA domains retain the ancestral switch function characteristic of GTPases as a whole.

Our analysis also revealed ParA branches with contexts indicating previously unstudied roles in core cellular functions. For instance, we found a gene-neighborhood coupling between PriB2-clade ParA genes and genes encoding HypA-domain proteins in certain archaea and bacteria (Figure 2B, panel 7). This suggests that in these systems, ParA has replaced another SIMIBI GTPase, HypB, as a potential metallochaperone delivering nickel to the [NiFe]-hydrogenase apoprotein [3, 94]. Further potential roles for two distinct PriB2-clade ParA NTPases in comparable processes are indicated by their two independent gene-neighborhood associations with the NifX and NifU domains, previously studied as accessory components of nitrogen fixation pathways (Figure 2B, panels 5, 6). One of these ParA clades contains an insert of a 4Fe-4S ferredoxin domain between the β2 and β3 strands (Supplementary Fig. S2A). These neighborhoods also feature additional genes, coding for a helix-turn-helix (HTH) domain, a protein with a poorly structured, glycine-rich region followed by a C-terminal highly charged helix, and sometimes additional 4Fe-4S ferredoxin domains, a metallo-β-lactamase domain, a Radical SAM metalloenzyme domain fused to a rapidly evolving winged HTH (wHTH) domain, and an ARMT1-like phosphatase (Supplementary Fig. S2A). This collection of associations with metalloenzymes and ferredoxins suggests a possible role for the system in metal-cofactor biogenesis, comparable to that of HypA-associated ParA proteins. Again, within the PriB2 clade, we found a noteworthy conserved gene neighborhood association between genes encoding ParA homolog and Type VII secretion systems (T7SSs; Figure 2B, panel 12), which, in Gram-positive bacteria, are involved in the secretion of protein toxins deployed in interbacterial competition [87]. We observed that ParA homologs in these T7SS contexts include both active and inactive versions with disruption of the catalytic residues. It is conceivable that they play a role in the subcellular localization of T7SS complexes.

We found that both the PriB1 and PriB2 clades contained separate monophyletic assemblages of ParA lineages with predicted immune roles (ConB1 and ConB2, respectively; Figure 2A). Both these assemblages display certain subtle distinguishing sequence features (Figure 2C). For example, ConB1 is marked by a highly conserved valine within Walker A, whereas ConB2 contains a conserved arginine in lieu of the typical Walker A lysine and a PG motif downstream of sensor-1 (Figure 2C). These observations suggest that the capacity of ParA proteins to act as switches in the assembly of macromolecular complexes has been repeatedly reused for immune functions, paralleling the situation with GTPases of the TRAFAC subgroup, such as dynamins and GIMAPs [15, 95]. These candidate ParA proteins involved in potential immune systems are described in below sections, and the results of multicellular enrichment test for these systems are shown in Figure 3.

**Figure 3.**
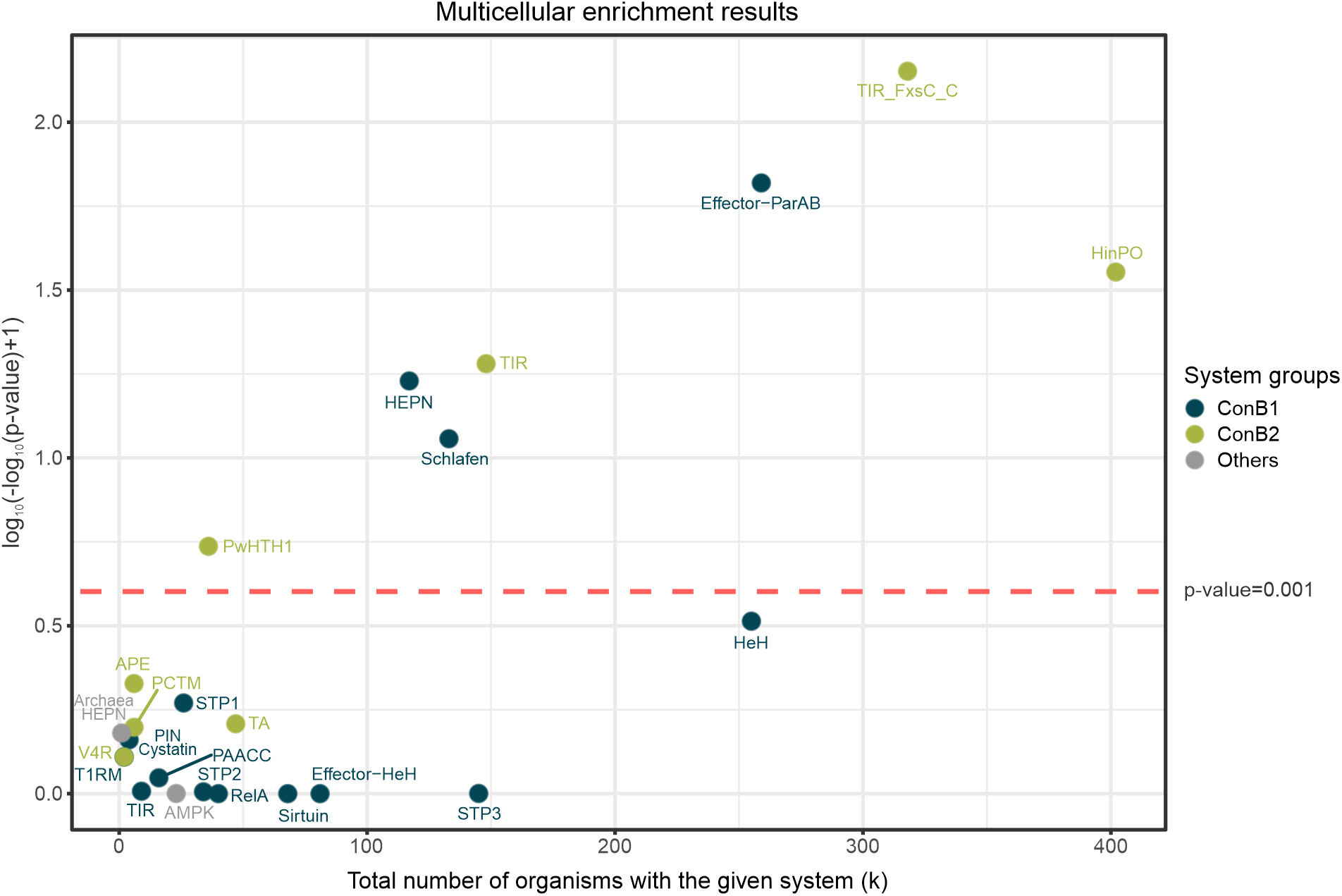
Multicellular enrichment results for prokaryotic ParA-based biological conflict systems. The x-axis shows the total number of organisms with the given system (k). The Y-axis shows values of log10(-log10(p-value)+1). A horizontal line at p-value=0.001 is indicated by a dashed red line. The legend for system groups is shown alongside. **Alt text**: Plot of multicellular enrichment for prokaryotic ParA-based biological conflict systems, showing system frequency versus statistical significance, with a dashed threshold indicating a p-value of 0.001.

### Systems with predicted immune roles in the ConB1 clade

#### The ParA-HeH system

A major contextual theme observed for a subset of the ConB1 assemblage couples ParA homologs in a two-gene neighborhood, with a gene encoding an uncharacterized, rapidly evolving protein. Through a combination of sequence profile and structural analyses, we found the protein encoded by the second gene to contain a previously unrecognized version of the Helix-extended-Helix (HeH) domain [96] (Supplementary Fig. S3). We designate these HeH domains as PHeH (for ParA-associated HeH) and these systems as ParA-HeH systems (Figure 4A, B). CLANS analysis of these domains revealed 22 distinct, highly divergent sequence clusters of PHeH domains (PHeH1-22). The AF3-predicted 3D structure of the ParA proteins from these systems revealed a characteristic highly variable insert region after core strand β2. Examination of the AF3-predicted structures of the PHeH proteins revealed that the HeH domain is linked to structurally variable α-helical N-terminal (and sometimes C-terminal) extensions (Supplementary Fig. S3). Hence, we posit that these variable extensions of the PHeH proteins may interact with the corresponding variable insert found in their partner ParA protein.

**Figure 4.**
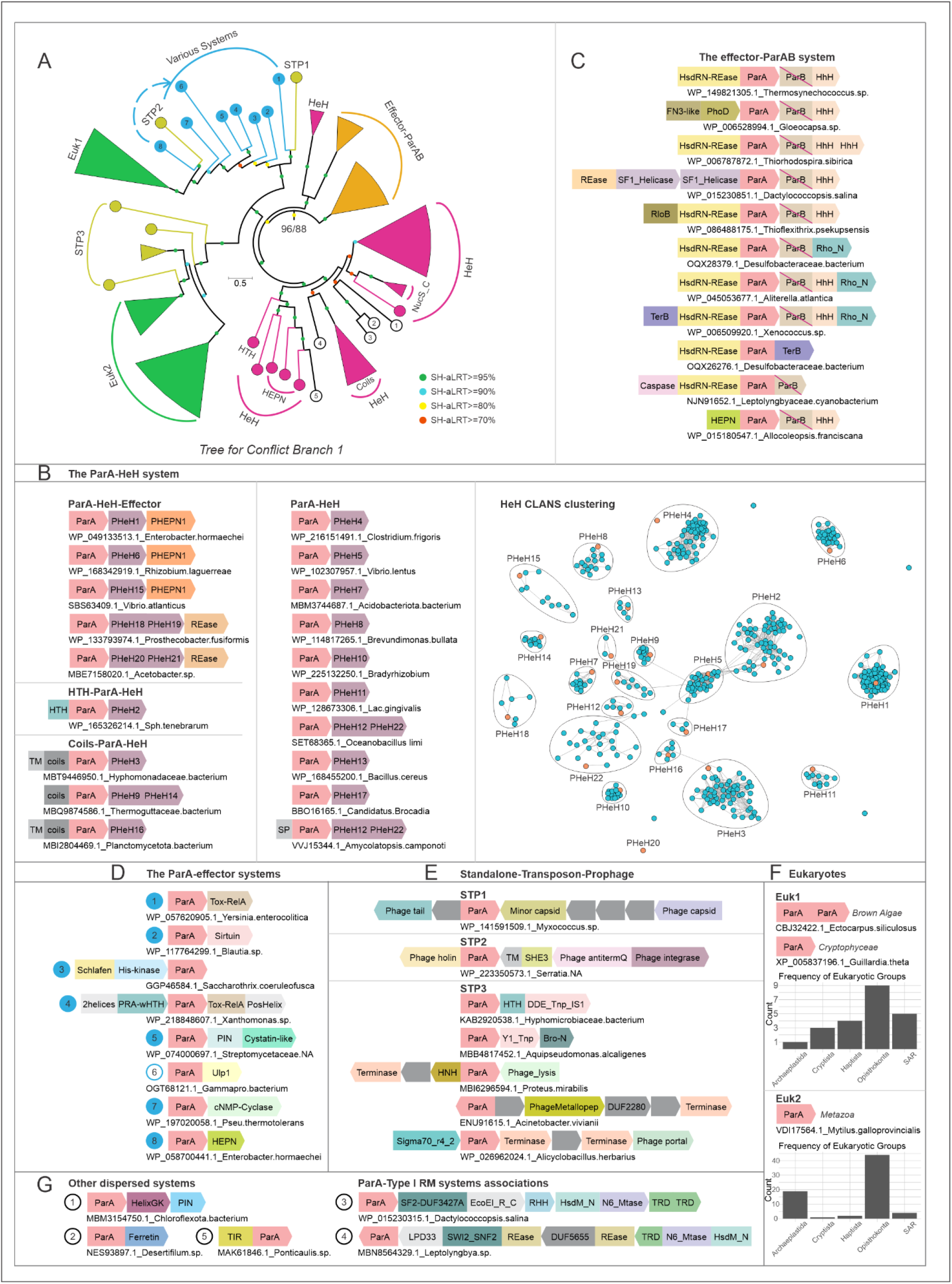
ConB1 (A) comprises diverse immune systems, including the ParA-HeH system (B), the effector-ParAB system (C), ParA-effectors systems (D), Solo-Transposon-Prophage (STP) clades (E), homologs transferred to eukaryotes (F), and several other dispersed systems (G). A CLANS analysis of PHeH domains from the ParA-HeH system illustrates their sequence diversity. A red across the domain indicates that it is predicted to be inactive. **Alt text**: ConB1 immune system diversity. Figure with subfigures labelled A to G, showing a phylogenetic overview of ConB1 immune systems, including the ParA-HeH system, the effector-ParAB system, ParA-effector systems, Solo-Transposon-Prophage clades, homologs transferred to eukaryotes, and other dispersed systems, with a CLANS analysis illustrating sequence diversity of PHeH domains.

We identified four subtypes of ParA-HeH systems based on the presence or absence of added components (Figure 4B): (i) the minimal subtype displays just the above-described two gene core; (ii) the subtype with an HTH domain fused to the N-terminus of the ParA domain; (iii) versions with a TM helix and coiled-coil region N-terminal to the ParA domain; (iv) the subtype with a third linked gene encoding an effector, such as a HEPN RNase (PHEPN1, Supplementary Fig. S2Q) or a NucS-related Restriction Endonuclease (REase) domain [97]. A representative of subtype 4 was recently characterized as an antiphage system and mediator of prophage competition in *Salmonella enterica*, with its HEPN domain catalyzing tRNA anticodon nuclease activity [98], indicating that our protocol indeed recovers genuine immune systems.

However, because a large fraction of these systems lacks the third gene coding for an effector nuclease, we posit that the basic action of these systems can be mediated solely by the ParA and PHeH components. Based on functional precedents of the conserved components, we propose that the PHeH domain acts as a sensor for the invading selfish element (a phage or a plasmid) by directly binding its DNA, akin to other HeH domains [96, 99]. Specifically, HeH domains might recognize specific DNA structural features, such as viral replication forks or chromosomal origins of replication. Additionally, the variable insert region in the ParA component and the extensions of the PHeH might also make contact with protein components of the invader element. This is supported by the high variability that we observed in all these regions, consistent with extreme sequence divergence driven by an arms race with the invader [100]. This recognition could then alleviate the system’s default negative regulation via downstream conformational changes. We suggest that the release of the PHeH-mediated negative regulation allows the ParA component to polymerize along chromosomes [35, 81, 101] and interfere with replication or segregation of the invader genome. Alternatively, (consistent with the derivation of ConB1 from classical ParAB systems; Figure 2A), the ParA could bind phage/plasmid-encoded ParB [35, 102, 103] and inhibit its function. The more elaborate subtypes outlined here (Figure 4B) likely augment this mode of action via the additional linked domains. For instance, the HTH domain fused to the ParA could provide a further nucleic-acid-interaction surface, while the TM and coiled coil segments could facilitate membrane localization and multimerization, respectively [28]. Finally, in versions with the third enzymatic effector, that component could serve as a second line of defense, which causes cell suicide in the event of the failure of the core system.

### The effector-ParAB system

The second major system featuring the ConB1 ParA domains is significantly overrepresented among bacterial taxa with a multicellular habit (e.g., cyanobacteria; see Figure 3 and Supplementary Fig. S1). It is organizationally related to the classical ParABS system in retaining the operonic ParA-ParB coupling. However, the residue conservation pattern of their ParB component indicates that it is likely enzymatically inactive (Supplementary Fig. S4). Nevertheless, it still retains several conserved polar residues in the vicinity of the canonical ParB active site, suggesting it may retain a nucleotide-binding capability comparable to the canonical version [104].

Further, both the ParA and ParB components of the effector-ParAB systems are distinguished by fusions to variable domains. The first variable domain is positioned immediately N-terminal to the ParA domain, and the second is C-terminal to the ParB domain (Figure 4C), in lieu of the constant HTH domain fused to the ParB in the housekeeping ParABS system [105]. The first variable position is occupied by a diverse range of enzymatic effector domains: while it is most commonly a Restriction Endonuclease superfamily (REase) domain, it also features the related calcineurin-like and PhoD-like phosphoesterases [31, 106, 107], RloB-like TOPRIM domain nucleases [108], Caspase peptidases [109], and HEPN RNases [110] (Figure 4C). In rare instances, the ParA is further fused to TerB domains, predicted to bind small-molecule ligands [111] (Figure 4C). The second variable position, in contrast, is occupied by one of several structurally distinct nucleic acid-binding modules, like the helix-hairpin-helix (HhH) [112] and Rho-N-type HeH domain.

Given the evolutionary relationship between the core components of this system and those of classical ParABS systems, we propose that they are likely to share a functional basis. In the classical ParABS systems, the ParB protein forms CTP-dependent dimeric clamps around chromosomal DNA and transiently interacts with ParA dimers that are non-specifically bound to DNA at some distance [113]. These tether-like interactions ultimately help promote chromosome partitioning [113]. In a similar vein, it is possible that in these systems, the ParB first recognizes specific DNA structures of viral or plasmid chromosomes via its variable C-terminal DNA-binding domain, and then forms a clamp around the targeted DNA by binding CTP or another nucleotide [35]. It may then recruit or associate with its cognate ParA on the invader chromosome. As the ParB in these systems is inactive, this complex is unlikely to dynamically associate and dissociate, and may instead prevent invader chromosome replication or transcription by coating the invader chromosome as a static polymer. It is also possible that this complex could interfere with the endogenous ParA and ParB of selfish elements that encode them. Finally, the N-terminal effectors fused to the ParA component could act as force multipliers or as a second line of defense, targeting cellular molecules to initiate programmed cell death or dormancy if the primary system fails. The latter is specifically supported by several of these effectors (e.g., Caspase, HEPN, REase) being known mediators of apoptosis in other defense systems [114–116]. More generally, the proposed role for these systems in interfering with invader chromosome dynamics is reminiscent of several antiviral defense systems featuring a P-loop ATPase of the ABC superfamily that function, at least in part, through manipulation of chromosomal organization [6, 117–121].

### The ParA-effector systems

Again in the ConB1 assemblage, we identified another branch displaying several potential immune mediators widely distributed across multiple prokaryotic lineages (Figure 4A; Supplementary Fig. S1). These systems couple a gene encoding a ParA protein with one coding for a variable enzymatic effector domain (Figure 4D). The coupled variable effectors include the following: (i) the Tox-RelA domain, which catalyzes nucleotide polyphosphate modification of target RNAs and probably other macromolecules [122, 123]; (ii) the Sirtuin domain, catalyzing NAD+-dependent reactions such as NADase [124], ADP-ribosyltransferase [125], or protein deacylase [126]; (iii) a Histidine Kinase fused to a Schlafen RNase domain [21, 127]; (iv) a PIN domain RNase [128] linked to a novel C-terminal domain adopting a Cystatin-like fold (Supplementary Fig. S2B); (v) a HEPN RNase domain [110]; (vi) an Ulp1**-**related papain-like peptidase domain [129]; (vii) a novel effector domain which we identified as a previously unrecognized cNMP-cyclase domain (hereinafter, ParA-Associated cNMP-Cyclase (PAACC) domain). Indeed, most of these domains have previously been observed as effectors in immune and other biological conflict systems [22, 26, 30, 130, 131]. The Schlafen and HEPN domain-associating ParA systems are specifically notable for their enrichment in prokaryotic taxa with a multicellular habit (Figure 3).

Given that in the classical ParABS system, ParA can bind DNA nonspecifically through its C-terminal extensions (αD–F) (Figure 1A, Supplementary Fig. S5) [81, 101], we propose that the ParA in this system directly intercepts invading viral or plasmid DNA via its cognate rapidly evolving C-terminal extensions. In several exemplars of this system, especially those with the Tox-RelA gene, there is an additional operonically coupled upstream gene encoding a rapidly evolving wHTH domain (Figure 4D). Interestingly, related wHTH genes are also frequently found as multiple genomically linked copies in a broader set of defense contexts (see the poly-wHTH systems below). These wHTH domains could augment the specificity of the ParA component of these systems towards viral or plasmid DNA. Thus, the ParA domains in these systems could potentially function as inhibitors that sequester invader DNA, preventing it from replicating or being transcribed. In the event this inhibition is overcome by invasive elements, the associated effectors are likely to be released, likely killing the host cell via a programmed cell death response specific to the effector’s biochemistry.

### Sporadic ParA lineages with potential immune functions

We observed three monophyletic lineages of ParA domains nested within the ConB1 assemblage, which are encoded within conjugative transposons or prophages (Figure 4A, E), leading us to designate them as the STP (Standalone-Transposon-Prophage) lineages. Furthermore, these STP lineages are sister groups to two eukaryotic clades of standalone ParA proteins (Euk1 and Euk2 in Figure 4A, F). These are primarily found in certain metazoans (with lineage-specific expansions of Euk2 in bivalve mollusks and rotifers), and photosynthetic eukaryotes with primary or secondary chloroplasts (Figure 4F). Phylogenetic analysis suggests that the eukaryotes likely acquired these versions via horizontal gene transfer (HGT) from bacterial STPs on two independent occasions (Figure 4A). We propose that the STP versions might act as a potential conflict system that targets rival phages or conjugative elements by interfering with their endogenous chromosomal segregation apparatus. In light of this, it is tempting to postulate that the eukaryotic versions might similarly defend genomes against selfish elements targeting them.

Within the ConB1 assemblage, several additional systems are sporadically dispersed within the ParA-HeH branch (Figure 4A, G). These systems feature contextual associations between ParA and the following: (i) a two-gene module encoding a PIN domain RNase and protein with a novel domain composed of helices and a Greek-key β-sheet (the HelixGK domain; Supplementary Fig. S2C); (ii) a NAD+-targeting TIR domain directly fused to the ParA [22, 132] reminiscent of the above-described ParA-effector systems; (iii) Type-I Restriction-Modification (R-M) systems (Figure 4G, panel 3, 4) [133]. The first system resembles the effector-encoding variant (subtype 4) of the ParA-HeH system, with the HeH module replaced by the HelixGK domain, suggesting that it might be functionally equivalent to the former. ParA domains embedded within Type I R-M systems are likely to function as single-component, second-line defenses that are activated when the R-M system fails, as has been described for other embedded effectors [134, 135].

### Systems with potential immune functions in the ConB2 clade

#### The PCTM system

In the ConB2 assemblage, a clade of ParA homologs is noteworthy for unusual receptor-like architectures: here, a cytoplasmic ParA domain is fused to a C-terminal transmembrane domain with a variable number of TM helices often followed by an extracellular/periplasmic region (a type-II single TM protein). This periplasmic region usually features one of several domains, such as pentapeptide repeats, TPRs, or a SPOR-like peptidoglycan domain (Figure 5A, B, Supplementary Fig. S2D). Accordingly, we designated the system centered on these proteins as PCTM (ParA with a C-terminal TM domain) (Figure 5B). The gene encoding this protein is coupled to an upstream gene that codes for a predicted effector domain. These effectors either feature classical domains (e.g., the SLATT, a putative 2TM effector or a Caspase) or the previously uncharacterized rapidly evolving, α-helical CPPC-4h domain, which is characterized by a CP-PC motif. In some cases, these systems contain a further downstream gene coding for an extracellular protein with a ligand-binding domain of the TAXI family [136]. We propose that these systems function by detecting the invader via intramembrane or periplasmic perturbations (e.g., of the peptidoglycan) caused by actions such as viral injection, capsid packaging or plasmid conjugation. We posit that these ParA domains function as a signaling switch that either activates the effector encoded by the upstream gene or triggers assembly of a membrane-linked multimeric complex. This action could limit the selfish element through programmed cell death or by inhibiting the assembly of its transmissive complexes (e.g., capsids or the conjugative apparatus).

**Figure 5.**
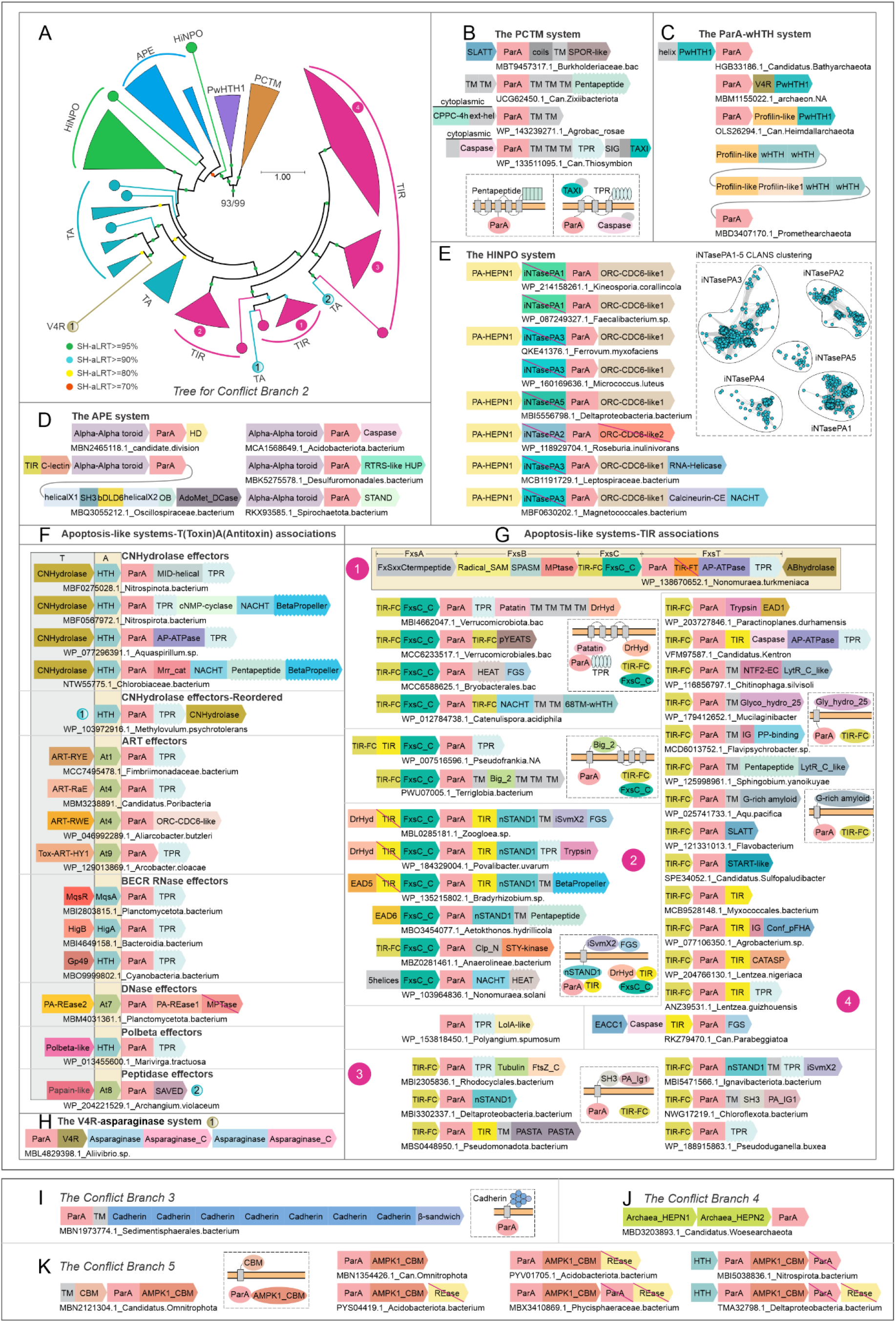
ConB2 (A) comprises diverse immune systems, including the PCTM system (B), the ParA-wHTH system (C), the APE system (D), the HinPO system (E), and apoptosis-like systems associated with toxin-antitoxin (TA) pairs (F) or TIR domains (G). Systems identified from ConB3 to ConB5 are shown below (I– K). Models of representative receptor variants are presented in dashed boxes. **Alt text**: ConB2 to ConB5 immune system diversity. Figure with subfigures labelled A to K, showing a phylogenetic overview of ConB2 immune systems, including the PCTM system, the ParA-wHTH system, the APE system, the HinPO system, and apoptosis-like systems associated with toxin-antitoxin pairs or TIR domains, with additional systems from ConB3 to ConB5 and models of representative receptor variants.

### The ParA-wHTH system

The second system in the ConB2 assemblage (Figure 5C), primarily of archaeal provenance (Supplementary Fig. S1), couples its ParA gene to one or more genes coding for a distinctive rapidly evolving wHTH DNA-binding domain (PwHTH1, Supplementary Fig. S2F), which is fused to an elongated N-terminal α-helix . We designate these systems the ParA-wHTH systems. In its simplest two-gene form, the system resembles the minimal version of the above-described ParA-HeH systems (subtype 1), with the wHTH acting as the potential DNA-recognition module in lieu of the HeH domain. In some cases, the wHTH component is elaborated via fusions to either a soluble-ligand-binding V4R domain [137, 138] or a previously uncharacterized domain (Profilin-like, Supplementary Fig. S2E) of the profilin-longin-like superfamily [139]. Interestingly, the V4R domain is also found directly fused to ParA in another branch of the ConB2 assemblage (Figure 5H). The profilin-longin-like domains frequently associate with TRAFAC and SIMIBI GTPases across the tree of life, including in other potential anti-selfish-element systems (HL and LA, unpublished observations) [140]. One of these is the previously described system [140], which combines a MglA GTPase with a MglB-like member of the profilin-longin-like superfamily [141] and a wHTH fused to an ABC-ATPase. Given the parallels to the ParA-HeH systems, we hypothesize that the ParA-wHTH systems function similarly to intercept and coat invader DNA via the combined action of the wHTH and ParA domains. The versions augmented by the V4R domain, which acts as a small-molecule-binding or an enzymatic domain[137, 138], could additionally respond to a soluble signal recognized by this domain. Versions elaborated with the profilin-longin-like domain could potentially extend their action in conjunction with the ParA protein through multimeric assemblies of that domain, as previously observed in eukaryotes [142].

### The APE system

This system, featuring a ParA protein of the ConB2 assemblage, spans several distant prokaryotic phyla (Supplementary Fig. S1). It is defined by a conserved three-gene neighborhood that brings together the ParA protein, a previously uncharacterized protein, and a variable candidate effector component. AF3 structure predictions identify an α-α-toroid domain at the N-terminus of the uncharacterized protein, followed by a long, charged C-terminal helix. Hence, we termed this system the APE (for <u>A</u>lpha-Alpha toroid-<u>P</u>arA-<u>E</u>ffector) system (Figure 5D). Interestingly, structure similarity searches revealed that the α-α-toroid domain is related to the thioester domain (TED) of human complement system proteins C3 and C4 (Supplementary Fig. S2G). The activation of C3 to C3b involves a structural rearrangement of the TED, exposing a thioester crosslink (Cys-Gln side-chain bond) that covalently attaches to the surface of target pathogens or cells [59, 143]. While the α-α-toroid domain in the APE systems lacks the conserved cysteine seen in the classical TED domain, it contains a conserved serine or threonine residue spatially positioned equivalent to the former residue. Hence, it is conceivable that this can form an ester linkage comparable to the C-Q thioester in TED. While it remains to be seen if this can form crosslinks akin to classical TED domains, we propose that the α-α-toroid domain, nevertheless, functions similarly as a sensor module for invading elements, while the C-terminal helix interacts with the ParA component, comparable to other tetherings observed between ParA and partner proteins [35, 113].

The variable effector component in the APE systems includes versions of the HD phosphoesterase [31], Caspase peptidase, TIR, and an NTPase related to the KAP NTPases [144]. In addition, we also recovered uncharacterized proteins in the effector position that have not been previously reported in antiviral defense systems. One of these proteins encodes an effector with a HUP superfamily Rossmannoid domain most closely related to the version found in class I aminoacyl tRNA synthetases (AATRS) and HIGH nucleotidyltransferases [145] (Supplementary Fig. S2H). Non-canonical AATRS-related enzymes have previously been reported to catalyze the formation of cyclic dipeptide antibacterial molecules [145, 146]. We propose that this domain may comparably synthesize modified peptide products with a potential anti-infection role. The second uncharacterized effector displays a core containing a central bacterial member of the Death-Like superfamily (bacterial Dealth-Like Domain 6: bDLD6) [29], flanked on both sides by helical domains followed in turn by a SH3-fold (N-terminus) and OB-fold domains (C-terminus) (Figure 5D, Supplementary Fig. S2I). This core may be further augmented by fusions to TIR and amino group acyltransferase (GNAT) domains. Systems containing this effector are typically accompanied by a further upstream gene coding for a TIR domain fused to either a C-type lectin or an inactive HKD domain (Figure 5D) and additional genes coding for a deiminase or AdoMet decarboxylase [147] (Figure 5D).

As with several other antiviral systems described above and in previous studies, we interpret the variable enzymatic effector component of the APE systems as a second-line defense that backs up the primary immune function provided by the constant Alpha-Alpha toroid-ParA pair. We hypothesize that the pair plays a role in forming a physical barrier or coat that limits viral infection, whereas the associated effector induces programmed cell death if the primary defense fails. In this regard, the presence of bDLD6 in a subset of these systems is particularly notable, as it points to the potential assembly of multimers of that protein through homotypic interactions, as has been reported for Death-like domains in metazoan programmed cell death systems [29]. This action could be extended by the associated SH3-fold and OB-fold domains, which could capture proteins or nucleic acids [148, 149]. The presence of the AdoMet decarboxylase is also reminiscent of other phage-host conflict systems, wherein it could deplete the key metabolite AdoMet required for several key processes, such as viral DNA modifications [150, 151]. The fusion of the TIR to the C-lectin domain in the auxiliary effector is again notable, as it might serve as an independent sensor or receptor for an invader signal (LM Iyer, AM Burroughs, L Aravind, personal observations).

### The HINPO system

This system, centered on a ParA protein from the ConB2 assemblage (Figure 5E), is striking for its significant enrichment in bacteria with a multicellular habit (Figure 3 and Supplementary Fig. S1). The ParA component is distinguished by a fusion to a rapidly evolving N-terminal domain. Using structure homology searches, we established that this domain belongs to the DNA polymerase β-like nucleotidyltransferase (Pol-β) superfamily [152], which has been previously observed in several distinct immune contexts: for example, SMODS, the CCA-adding enzyme-related domain, and the minimal nucleotidyltransferase and RelA/SpoT [22, 31, 153] (Supplementary Fig. S6). However, this version is distinct from those versions, and an inspection of its alignment revealed that it is catalytically inactive (hence, inactive nucleotidyltransferase: iNTase). Here, the ParA component is sandwiched in a conserved gene-neighborhood by an upstream gene for a previously uncharacterized HEPN domain (PA-HEPN1 for ParA-associated HEPN 1) (Supplementary Fig. S2Q), and a downstream gene coding for a ORC-CDC6/ STAND clade AAA+ NTPase protein [4] (Supplementary Fig. S7). In several cases, the ParA component with its iNTase domain is directly fused to the ORC-CDC6/STAND-like component. We accordingly designate the systems the HINPO (<u>H</u>EPN-Inactive <u>N</u>ucleotidyltransferase-<u>P</u>arA-<u>O</u>RC/CDC6) systems (Figure 5E).

ORC-CDC6 and other AAA+-NTPases of this clade, like the transposase ATPase subunits, are known to bind DNA, including replication origin sites and transposon termini [154, 155]. Hence, we posit that the ORC-CDC6 in these systems acts as a sensor for selfish-element DNA and guides the formation of multimeric assemblies of the ParA component to block infection. Indeed, this mode of action is analogous to that proposed for the effector-ParAB systems from the ConB1 assemblage (Figure 5E). Given this proposed mode of action, comparable to the effector-ParAB and the HINPO systems described above, the PA-HEPN1 component would act as the backup effector, inducing cell death or a dormancy response [134]. The more enigmatic element of these systems is the iNTase domain: structural modeling suggests it retains the ability to bind a substrate (Supplementary Fig. S6); thus, it could act as a sensor that binds nucleotides or nucleic acid fragments. This raises the possibility that these systems are under dual regulation, with the ORC-CDC6-like component sensing DNA and the iNTase component sensing diffusible nucleotides or degradation products of nucleic acids. While the sources of these nucleotides remain unclear, they could be generated by phage-encoded metabolism, degradation of host nucleic acids by viral restriction attack or signals generated by other immune systems with nucleotide synthetases.

### Pervasive presence of ParA in predicted apoptosis-related systems

A widespread feature of the previously discussed systems is the presence of effectors, which we interpret as inducing cell suicide or dormancy responses to back up a core ParA-centered mechanism. In contrast, we also found a striking array of systems with a ParA protein from the ConB2 assemblage, wherein an apoptotic response appears to be their primary mode of action. The first class of these systems recruits diverse toxin-antitoxin (TA) systems as their effector module. The second class is defined by the presence of a TIR domain component and shares several domains and hallmark features of eukaryotic apoptosis systems. We consider these separately in the sections below.

### ParA-based apoptosis systems that have captured a Toxin-Antitoxin unit

Type II TA systems typically feature a toxin under the negative regulation of a transcriptional repressor antitoxin [156]. The systems under consideration are extensions of type II TA systems with a third component, namely a ParA domain fused to one of several variable C-terminal sensor domains (Figure 5F, Supplementary Fig. S8A). While this C-terminal sensor domain is most commonly TPRs, it might include nucleotide-sensing SAVED and inactive MPTase domains [22, 157] or a whole module centered on an NACHT family STAND NTPase fused to a C-terminal β-propeller domain [30]. Notably, we observed that the gene for the ParA component occasionally inserted between those coding for the toxin and antitoxin components (Figure 5F). This supports a scheme wherein the products of the three genes form a ternary protein complex. While three-component variants of type II TA systems have been described with a SecB chaperone domain protein that assists in folding of the disordered antitoxin [158, 159], the domain architecture of the ParA component makes these an entirely distinct theme. Instead, their organization is reminiscent of multiple recently described prokaryotic immunity-linked apoptosis systems that have a comparable ternary organization [28, 29].

These ternary ParA-containing systems can be classified into at least 6 subtypes based on their toxin domain (Figure 5F): (i) *carbon-nitrogen hydrolase (C-N hydrolase) toxins*. These are typically associated with an HTH antitoxin. Known C-N hydrolase domains catalyze the hydrolysis of nonpeptide carbon-nitrogen bonds [160]; hence, we propose that these could deplete essential cellular metabolites, like nucleotides, or cleave the nucleobase-sugar linkage in nucleic acids. (ii) *ADP-ribosyltransferase (ART) toxins* (Supplementary Fig. S2J), which modify target molecules with ADP ribose (ADPr) derived from NAD+ [161]. Some members of this superfamily catalyze a similar reaction to degrade NAD+ by cleaving off ADPr [162]. Accordingly, these toxin domains likely act either as ADP-ribosylating or NAD+-depleting toxins, as previously described in numerous conflict systems [163, 164]. These are coupled to several structurally distinct, previously unrecognized antitoxin domains. (iii) *Barnase-EndoU-ColicinE5/D-RelE (BECR) RNase superfamily toxins* [87]. These include a diversity of TA pairs unified by an RNA-cleaving toxin of the BECR superfamily coupled with an HTH antitoxin, such as MqsR-MqsA [165], HigB-HigA [166], and Gp49-HTH [167]. Previously characterized members (e.g., MqsR and HigB) have been shown to cleave mRNA to shut down translation. (iv) *REase superfamily toxins* (Supplementary Fig. S2K). These are predicted to cleave genomic DNA [116, 168]. (v) *Pol-β nucleotidyltransferase toxins* (Supplementary Fig. S2L). These might evince their toxicity by modifying target macromolecules with nucleotide adducts, comparable to the MenA1 and MenA3 toxins, which add pyrimidines to tRNA acceptor stems to inhibit translation [169, 170], or by generating a soluble nucleotide signal [21]. (vi) *Papain-like peptidase toxins* (Supplementary Fig. S2M). These are predicted to cleave target proteins upon induction.

### The TIR-ParA apoptosis systems and their diverse ramifications

These systems are defined by conserved gene neighborhoods that couple the ParA gene to an upstream gene encoding a conventional TIR domain, with a NAD+-degrading active site (Figure 5G, Supplementary Fig. S8B). In a subset of these systems, the ParA domain is fused to a further C-terminal TIR domain that appears to have lost its NAD+-degrading active site. In one example of this class of systems from Actinomycetota, the active TIR component maps to the FxsC protein, and the ParA, with the inactive TIR fused to it, maps to the FxsT protein (Figure 5G, panel 1). These were named thus for their genomic clustering with the FxsAB peptide-processing system, but remain functionally uncharacterized [171]. Accordingly, we hereafter distinguish the conserved active TIR domain as TIR-FC (TIR in <u>F</u>xs<u>C</u>) and the inactive version fused to the ParA as TIR-FT (TIR in <u>F</u>xs<u>T</u>). The TIR-FC domain is often fused to a C-terminal FxsC-C domain, which we establish to be an additional inactive TIR domain (Figure 5G, Supplementary Fig. S2N). Another hallmark of these systems is the architecture of the ParA component: irrespective of whether the ParA domain is fused to a C-terminal TIR-FT or not, it shows fusion to a wide range of additional domains, several of which are found as potential sensors of invader presence in other immune contexts (Figure 5G; see below).

These systems can be classified into several subtypes based on the architectural theme evinced by the ParA component. We outline these below:

*(i) Fusions to STAND NTPases*. A major subtype (Figure 5G, panel 1, 2) has the ParA module ‘grafted’ onto the N-terminus of one of several STAND NTPase modules (AP-ATPase, nSTAND1, and NACHT) previously described in immunity and apoptosis pathways in both prokaryotes and eukaryotes [4, 29, 30]. The STAND module has a characteristic tripartite architecture with an N-terminal effector domain (e.g., TIR or Caspase) followed by a central AAA+ NTPase and C-terminal superstructure-forming repeat domains (e.g., TPR, HEAT, pentapeptide, or β-propellers). STAND modules sense the presence of invaders via the C-terminal superstructure-forming domain and trigger apoptosis by activating the N-terminal effector [30].
*(ii) Ligand-binding subtype*. Here, the ParA component is linked to a C-terminal ligand-binding domain, including several previously reported in biological conflict contexts, namely pYEATS [21], FGS [28], and CATASP [21]. In addition to these, we also identified sensor domains previously unrecognized as functioning in immune contexts. Two of these, belonging to the START [172] and FHA [173] superfamilies, define novel, widespread anti-selfish element defense strategies independently of their ParA association (see below) (Figures 5G,6C-D). A third new sensor domain was unified with the LolA superfamily, which binds lipid ligands in the periplasm [174, 175]. We predict these systems respond to viral presence by sensing soluble metabolites that might change in concentration during infection or specific phage-induced ligands.
*(iii) Receptor-like subtype*. Here, the ParA domain is the constant N-terminal cytoplasmic domain of a type-II single-transmembrane protein with a variable C-terminal extracellular sensor domain analogous to the version described above in the PCTM systems (Figure 5G, panels 3 and 4; Supplementary Fig. S2O and S2P). The extracellular sensor can take the form of peptidoglycan-binding domains (e.g., PASTA, LytR-C, Ig-like, and bacterial SH3) [176–178], domains that likely process peptidoglycan (e.g., TIM-barrel fold glycosyl hydrolase family 25) [179], or domains predicted to bind to other ligands (e.g., Phosphopantetheine-binding superfamily) [180]. We also identified a previously uncharacterized glycine-rich extracellular domain in these systems that is predicted to adopt an amyloid structure with potential adhesive surfaces. This amyloid domain occurs outside the ParA association in other secreted bacterial proteins fused to peptidoglycan-binding domains, such as SPOR and OmpA, as well as in a novel caspase-based conflict system (LM Iyer, AMB, LA, unpublished observations). These systems are predicted to intercept invasion signals (e.g., the breaching of peptidoglycan) in the periplasmic/extracellular space [181].

**Figure 6.**
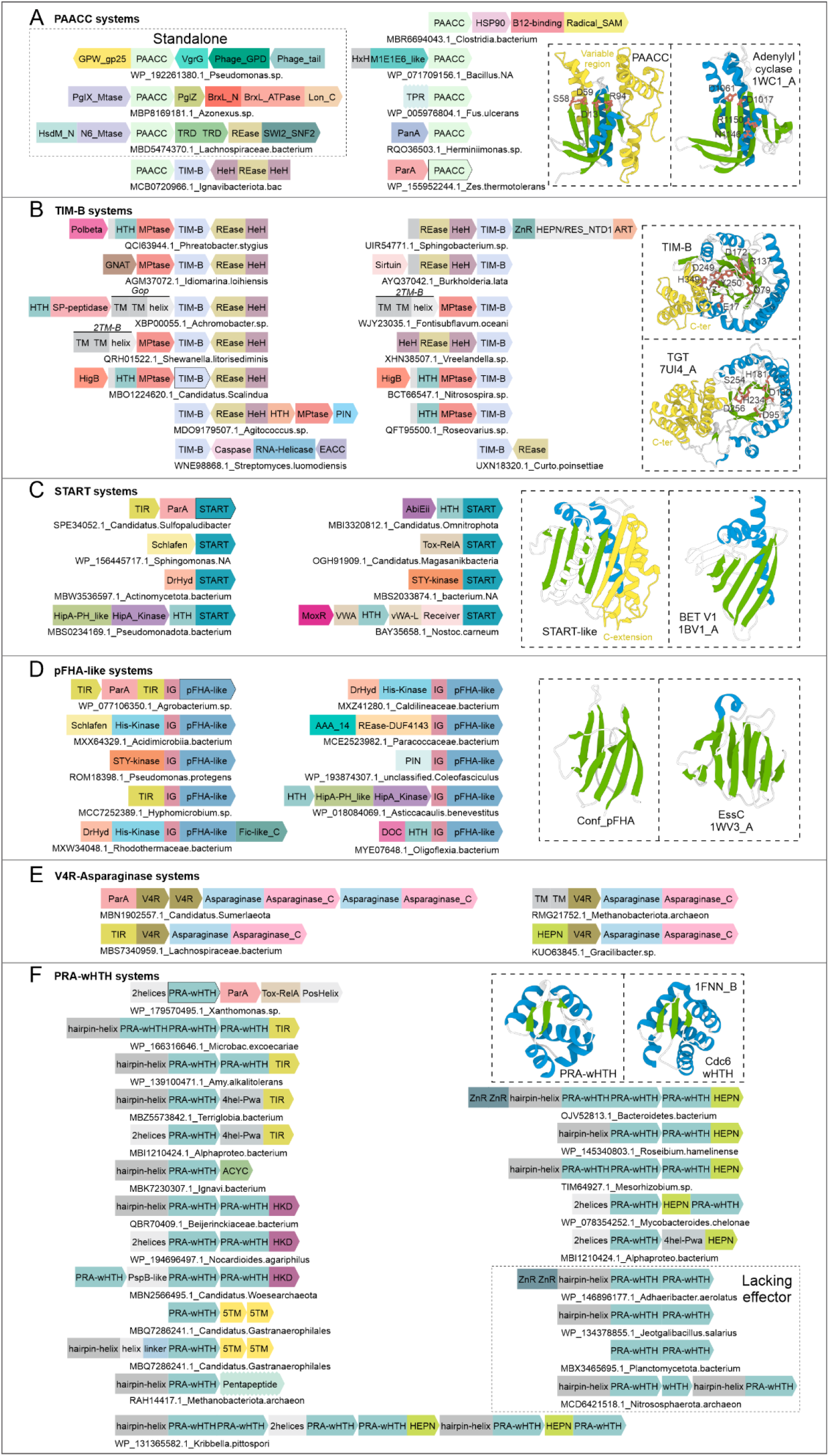
Predicted immune systems based on domains originally identified in association with ParA: PAACC-based systems (A), TIM-B-based systems (B), START-based systems (C), pFHA-like-based systems (D), V4R-asparaginase-based systems (E), and PRA-wHTH-based systems (F). Cartoon representations of these domains are shown, and conserved active sites are depicted as ball-and-stick models for enzymatic ones. For the newly identified domains (PAACC, TIM-B, START, pFHA-like, and PRA-wHTH), homologs identified by structural comparison are also shown. **Alt text**: Predicted immune systems from ParA-associated domains. Figure with subfigures labelled A to F, showing predicted immune systems based on domains originally identified with ParA, including PAACC-based, TIM-B-based, START-based, pFHA-like-based, V4R-asparaginase-based, and PRA-wHTH-based systems, with cartoon representations of the domains and conserved active sites shown for enzymatic ones.

Beyond these widespread architectural subtypes, rarer variations are observed in these TIR-ParA systems (Figure 5G, Supplementary Fig. S9). The include: (i) versions where the constant TIR domain (TIR-FC) component is N-terminally fused to or displaced by either effector-recruiting homotypic adaptor domains of the Death-like superfamily (e.g., EAD5, EAD6) [29], or other effector domains (e.g., DrHyd, Trypsin, or a Caspase as part of the previously described EACC1 system [28]). (ii) Versions which lose the ParA domain entirely, with a direct fusion of the TIR-FC domain to a STAND NTPase module. (iii) Variants where the TIR-ParA gene-dyad is incorporated into a larger mobile gene-neighborhood, which codes for a modified peptide-processing system typified by FxsAB (Figure 5G and Supplementary Fig. S8) [171].

### Functional models for ParA-centric apoptosis-related systems

While the two types of systems described above differ in their components beyond ParA, they still share a basic organizational logic. This is reflected in the strong conservation of gene order across operons of these systems, suggesting that they form a multimeric complex whose spatial organization is critical for their function. In both cases, it appears that the ParA component acts as a switch that regulates the initiation of cell death in response to a signal of invader presence sensed by the fused sensor domains. The diversity and subcellular localization of these sensor modules further suggest that they have been selected to recognize a wide range of signals that might either be directly derived from the invader or be secondary metabolic consequences of invasion. Moreover, these signals might be intercepted both within and outside the cell. However, beyond this shared process, the two types of systems likely differ in both their deployment contexts and modes of action. An important clue in this regard comes from the observation that, in contrast to TA-ParA systems, TIR-ParA systems are significantly enriched in bacteria with a multicellular habit, particularly when the FxsC-C domain is present (Figure 3 and Supplementary Fig. S1). Thus, the former systems appear to be a regulated extension of the pan-prokaryotic Type-II TA principle, which has been recruited for a potential antiviral role by the accretion of the ParA-sensor third component. Conversely, the TIR-ParA systems appear to be specifically geared to act in multicellular contexts. Hence, we suggest that they are systems primarily aimed at limiting viral spread within a multicellular ensemble via the “self-sacrifice” of individual infected cells by apoptosis.

For the TA-ParA systems, given their ternary organization, we postulate that under default conditions, the toxin component is negatively regulated by its association with the antitoxin. However, upon the sensing of the presence of a selfish element by the sensor modules, the associated ParA domain is posited to act as a switch reorganizing the toxin-antitoxin interaction and unleashing the toxin to induce apoptosis [18, 182]. The mode of action of the TIR-ParA systems, while broadly similar, can be best understood as a cascade. Here, sensing of the signal indicating invader presence by the sensor domains is likely transmitted to the associated ParA domain, which then acts as a switch, triggering the active TIR component to catalyze the destruction of NAD+ stores in the infected cell. In systems with a STAND NTPase module, activation is likely a two-step cascade, with the STAND NTPase triggering the first NTP-dependent step—potentially via multimerization—followed by the second NTP-dependent step mediated by the ParA domain. Such a multistep process could help set a threshold for activation of lethal effectors and limit their deployment by “false alarms.” The action of these systems is further complicated by the presence of one or more catalytically inactive TIR domains (TIR-FT and FxsC-C). While they are unlikely to degrade NAD+, like other TIR domains, they might still participate in multimerization, supporting the idea that these systems might form large multimeric assemblages as reported for other apoptotic systems [183, 184]. Finally, the association of a subset of these systems in Actinomycetota with the FxsAB peptide modification and processing system suggests that their apoptotic action could be coupled with the release of a modified peptide that acts either as an antiviral molecule itself or a diffusible signal “alerting” adjacent cells in the multicellular assemblage to the infection [28].

### Other sporadic ParA lineages with predicted immune roles

Several smaller ParA lineages evincing characteristics of potential immune systems were observed outside of the ConB1 and ConB2 assemblages described above (Figure 2A).

### ParA domains in standalone receptor-like proteins

The domain architecture of these systems from the *Phycisphaerae* phylum is thematically equivalent to the above-described single TM receptor-like architectures found in the PCTM and third subtype of TIR-ParA systems (Figure 5I). However, they lack the other associations of the above systems, and the ParA belongs to a distinct branch of the tree (ConB3 in Figure 2A). Here, the intracellular ParA domain is fused to a TM helix and a C-terminal extracellular/periplasmic region displaying a tandem series of cadherin domains capped by a distinct C-terminal β-sandwich domain. Their organization suggests that they sense infection-induced alterations to cell-surface carbohydrates in a calcium-dependent manner to transduce a defense signal via their intracellular ParA domain [185].

### Carbohydrate-binding-module (CBM)-containing system

In this system, sporadically distributed across several bacterial phyla, the ParA domain is fused to a C-terminal β-jelly-roll fold lectin domain (CBM; Pfam: AMPK1_CBM), which is known to bind polysaccharides such as glycogen [186] (ConB5 in Figure 2A and Figure 5K). The rapid divergence of the CBM domain points to a potential role for these systems in immune function. This core system might be elaborated by two alternative extensions. The first of these adds a further upstream gene encoding a single TM protein with a second, fast-evolving extracellular/periplasmic CBM domain (Figure 5K). In the second variant, the core ParA-CBM gene is joined by genes encoding a second rapidly evolving ParA and REase proteins, both of which are predicted to be catalytically inactive. Given that the CBM lectin domain is the defining feature of these systems, they likely function by detecting carbohydrates, triggering ParA multimerization to limit infection. We had earlier proposed that a wide range of lectin domains have been recruited as immune sensors in both bacterial and eukaryotic antiviral systems [29]. They could sense infection by either directly binding glycosylated virion proteins or carbohydrate products generated as a consequence of infection, such as peptidoglycan fragments or capsular precursors induced by phage infection. They could also detect polysaccharide depletion, which has been reported to be induced by cyanobacterial phages [187]. This basic sensory mode could be further extended in some of these systems by the inactive REase domain detecting other signals, such as DNA.

### Archaeal dual HEPN system

This is restricted to the archaeal clade of Nanobdellati and combines the ParA gene with two genes coding for previously uncharacterized HEPN domains (ConB4 in Figure 2A, Figure 5J and Supplementary Fig. S2Q). The HEPN domains are rapidly diverging at the sequence level (Supplementary Data), even between closely related species, suggesting rapid evolution in an arms-race scenario due to their functioning in an immune context. Further, an examination of the conservation pattern of these HEPN domains reveals a loss of the catalytic residues characteristic of this superfamily [110]. Hence, they are likely to be inactive as RNases while potentially still binding RNA (Supplementary Data). Given these observations, we propose that this system might recognize selfish elements by binding RNA targets, triggering the polymerization of the associated ParA to limit infection progression.

### Newly identified domains help identify novel, ParA-independent conflict systems

Due to the modular genomic architecture of conflict systems with a tendency to evolve via the recombination of basic functional modules, newly characterized domains can lead to the discovery of additional systems through analysis of the totality of their gene-neighborhood and architectural contexts [49, 87]. Accordingly, we analyzed the newly recovered domains in the above ParA-based conflict systems to identify additional systems where they occur independently of a ParA component.

### PAACC systems

We first identified the “ParA-Associated cNMP-Cyclase” (PAACC) to be associated with the ParA domain from the ConB1 assemblage (Figure 2A,4D). Consistent with a role in immunity, we found that despite being an enzymatic domain with intact catalytic residues (Figure 6A, Supplementary Data), PAACC is rapidly evolving, indicative of being locked in an arms race with selfish elements. We observed that PAACC also exhibits contextual associations independent of ParA, supporting its immune role. For instance, it is occasionally embedded within operons of other bacterial immune systems, namely R-M and PglZ/Brex (Figure 6A), in a manner reminiscent of several backup effectors [153, 188, 189]. This suggests that it is a potential effector that can be unleashed if the primary system, such as R-M, fails to contain an infection.

Other PAACC genes are part of several distinct conserved gene neighborhoods (Figure 6A), namely: (i) In a two-gene dyad with a gene encoding a PanA domain that has been found elsewhere in TA systems as an antitoxin inhibiting diverse associated toxins [190]. (ii) In an operon with a gene for the TIM-B protein (Pfam: PF14350, ‘Beta_protein’), which we unify here with the TIM-barrel fold. The founding member of this family is the phage P4 ‘B’ protein (hence, TIM-B from <u>TIM</u> barrel <u>B</u>), which negatively regulates the genomically linked Gop toxin [191] (see below). (iii) As part of a gene dyad with a gene encoding a two-domain protein with an M1E1E6 domain (a novel domain consisting of a four-stranded β-meander followed by three helices stacked onto it) and another previously uncharacterized helical domain with a conserved HxH motif. (iv) A three-gene neighborhood additionally encoding an HSP90 chaperone and a Radical SAM enzyme. (v) In phage genomes in the phage tail assembly operon. Given that both the TIM-B and PanA domains have been previously characterized as negatively regulating the activity of associated toxins [190, 191], we interpret these contexts as TA-like systems where the effector activity of the PAACC component is inhibited by its partners under default conditions, only to be unleashed during an infection. Analysis of the contexts of the M1E1E6-like and HxH-motif domain module suggests a similar antitoxin-like role. These observations support an effector role for the PAACC domain through its enzymatic activity, generating a toxic nucleotide or nucleotide modification of a target molecule

### TIM-B systems

We first uncovered the TIM-B domain as part of the above-described two-gene system in which it is coupled to PAACC. AF3 modeling followed by structural homology searches revealed that the TIM-B domain is specifically related to the TIM barrel domain of the deazaguanine transglycosylase family, which includes the Queuine tRNA transglycosylase (QTG), the DNA deazaguanine insertase (DpdA) and the enzyme modifying the N-terminus of the SMODS enzyme with a deazaguanine base [192–194]. TIM-B shares the diagnostic C-terminal hairpin and helix extension, which, in the deazaguanine transglycosylase family, positions the macromolecule substrate for deazaguanine insertion (Figure 6B, inset). Given that TIM-B possesses conserved residues that are spatially equivalent to the active site of the deazaguanine transglycosylase family [195, 196] (Figure 6B), it is predicted to catalyze a reaction involving a nucleobase transfer.

A comprehensive analysis revealed that TIM-B is coupled to a diverse set of alternative effector genes in addition to PAACC (noted above) or the P4 phage Gop toxin, which we predict functions as a 2TM membrane-permeabilizing effector. These effectors encompass a diverse array of biochemistry (Figure 6B): a BECR RNase (HigB), a previously uncharacterized REase (see below), the Pol-β effector described above in ParA contexts, an amino-group acyltransferase (GNAT), a sirtuin, an ART toxin of the RES family, a Caspase peptidase system and a second family of 2TM domains (2TM-B) distinct from Gop. Layered onto this core framework, a gene encoding a metallopeptidase domain, sometimes further fused to N-terminal HTH domains, is often found between the effector and TIM-B components or as an alternative effector (Figure 6B). In the REase effector, the endonuclease domain is fused to helical extensions at both termini, including a conserved C-terminal HeH domain (Figure 6B). In versions prototyped by the bacteriophage P4 system, there appear to be two effector components: TIM-B and the associated effector gene in the same orientation are linked to the said REase encoded by an overlapping gene transcribed convergently from the opposite strand. This organization of the REase gene was found to regulate the Gop-TIM-B operon, potentially at the transcript level [191].

The above observations, together with the prior results from the P4 system [191], suggest that TIM-B acts as a constitutive negative regulator of the associated effector via its deazaguanine transferase-related activity, either by generating an inhibitory soluble product or a covalent modification of the effector. Indeed, such regulation of antiphage response has been recently described for cyclic nucleotide generated by the mCpol system [197] and covalent modification of SMODS by the deazaguanine transferase [192]. The system could then be activated via disruption of the negative regulation by an infection marker, which might include phage proteins that sequester the TIM-B product or its substrates. In more elaborate systems with an MPTase component, release of TIM-B repression could be followed by an additional regulatory step involving proteolytic derepression reminiscent of the SOS and DdrO-IrrE systems [31, 198, 199]. Finally, in versions with a convergently transcribed REase gene, the transcript level regulation via potential mutual antagonism during transcription from overlapping opposite strands could result in alternative expression of either the REase or the TIM-B-effector pair, with one being unleashed if the other is suppressed by selfish-element countermeasures.

### START domain-based systems

We identified a previously uncharacterized version of the START domain superfamily in the sensor position of the above-described TIR-ParA systems (Figure 5G). This START domain is distinguished from all other previously characterized members of the superfamily by a C-terminal extension that contributes two additional strands to the core β-sheet (Figure 6C, inset). We further observed this type of START domain outside of the TIR-ParA systems coupled in an operon or directly fused to biochemically diverse predicted effector domains previously reported in other biological conflict systems, including: (i) those that degrade NAD+ (TIR, DrHyd); (ii) nucleotidyltransferases (AbiEii [200], and Tox-RelA [123]); (iii) RNases(Schlafen [130]); (iv) protein kinases (HipA [201] and STY-kinase); (v) HTH domains which could bind nucleic acids. Further, we found rare examples where it is incorporated into the previously described MoxR-vWA ternary systems predicted to induce apoptosis in multicellular bacteria [28]. START domains are known to bind lipid head-group ligands, such as sterols and phosphatidylcholine [172], and to regulate the activities of co-occurring domains in multidomain proteins [202]. Accordingly, we predict that these are defense systems in which the START domain senses alterations in the membrane caused by infection, triggering the associated effector domain, which can induce programmed cell death.

### Permuted FHA (pFHA) domain-based systems

We again originally recovered the pFHA domains as a potential ligand-binding domain in the TIR-ParA systems. Further analysis revealed that it is structurally comparable to the permuted FHA domains found at the N-terminus of the EssC protein of the Type VII secretion systems (Figure 6D) [74]; however, it is not closely related to those in sequence. Moreover, the pFHA domain does not conserve the phosphopeptide-binding histidine residue of classical FHA domains. We found an obligate association of this domain with an N-terminal Ig-like β-sandwich domain. Contextual analysis revealed that this two-domain module is further fused to the same array of biochemically diverse N-terminal effector domains as those reported above for the novel START domain, with some additional ones such as a PIN RNase, an REase (DUF4143-related), and a DOC [203] domain. We posit that these are likely to function similarly to the START domain systems, but by recognizing a distinct set of ligands, perhaps peptides rather than lipid head groups.

### V4R domains and asparaginase-based conflict systems

While the V4R domain has been observed across prokaryotic life as a ligand-binding or enzymatic (aliphatic reductase) domain [137], this work suggested that versions of it fused to an N-terminal ParA domain might be involved in immune functions (see above, Figures 5H,6E). Further analysis of related V4R domains revealed that the ParA domain can be replaced by a range of effector domains, including TIR, HEPN, and a predicted membrane-permeabilizing 2TM domain (Figure 6E). ParA occupying an equivalent position as these effector domains provides further evidence that ParA can itself act as an effector in this context. As with the above-described ParA-containing systems (Figure 5H), these systems are coupled to an additional gene encoding an asparaginase-family enzyme (Figure 6E). We propose that sensing of infection by these systems triggers the asparaginase-family enzyme to generate a diffusible messenger from asparagine or some other amide-containing substrate, as has been reported for some asparaginase-family enzymes [204, 205]. This ligand then likely binds the V4R domain to activate the fused effector domain.

### Poly-wHTH systems

In the ConB1 lineage, we identified a class of systems defined by the coupling of ParA with diverse effectors (Figure 4D). In these systems, when the effector is Tox-RelA, we identified an additional upstream gene coding for a previously uncharacterized and rapidly evolving wHTH domain (Figure 6F, inset). Further inspection of this wHTH domain revealed several parallel associations, wherein the ParA-Tox-RelA gene pair is replaced by several alternative downstream genes coding for other effectors (most commonly HEPN but also a cNMP-cyclase, HKD phosphoesterase, and TIR) across diverse prokaryotic taxa (Figure 6F). Several of these systems are notable for the presence of multiple wHTH domains, either as distinct genes or as tandem copies in the same polypeptide, with some systems featuring six or more wHTH copies (Figure 6F). Accordingly, we label these as poly-HTH systems. A subset of these systems contains only the run of wHTH domains, without any obvious downstream effector. Further structural variability is observed at the N-terminus of these wHTH proteins, with one or more of at least four distinct extensions (Figure 6F). The rapid sequence diversification of these wHTH arrays is consistent with selective pressure to co-evolve with a comparably diverging target, likely the genome of a selfish element. Thus, the primary action of these systems could be to coat invasive nucleic acids, thereby limiting their expression or replication. Under this framework, the associated effectors, when present, could serve as a backup to the primary response.

## Discussion

### Origin of the ParA-like domains

The SIMIBI subgroup of GTPases had already functionally diversified prior to the Last Universal Common Ancestor (LUCA) of Life [3] into lineages such as SR and SRP54, associated with the secretion machinery [206], and Mrp/NBP35, mediating iron-sulfur cluster assembly [207]. Importantly, the Mrp/NBP35-like clade diverged from the ancestral SIMIBI GTPases through degradation of the [NT]KxD motif and a concomitant change from GTP to ATP specificity. The two deep lineages of the ParA-like clade, viz., ParA proper and MinD, emerged from Mrp/NBP35 in the stem of the bacterial superkingdom, likely driven by the increasing size of chromosomes and were recruited to roles in chromosome segregation and cell division [3].

### Expanding the functional scope of ParA-like domains

In this work, we systematically explored the evolutionary trajectories of the ParA-like clade of SIMIBI GTPases (Figure 2A) and expanded their functional breadth (Figure 2B). We noted that some members of the ParA-like clade have potentially atavistically reacquired roles in the biogenesis of metal clusters, in addition to a plethora of roles in the subcellular organization of macromolecular complexes, and as switches comparable to GTPases from the TRAFAC subgroup (e.g., displacement of MglB, Figure 2B, panel 16). Here, we extended preliminary observations [28] to show that the ParA-like clade of the SIMIBI subgroup has also been repeatedly recruited for immune functions. Strikingly, our contextual analysis predicted that the ParA-like domains interface with an extraordinary range of partner components in these immune roles (Figures 4-5). These systems have repeatedly drawn on multiple distinct ParA-like lineages, indicating that these NTPase domains are eminently suited or “pre-adapted” for roles in anti-selfish element defense (Figure 2A). Thus, the ParA-like clade of the SIMIBI subgroup joins GTPases of the TRAFAC subgroup, including GIMAPs, the DO-GTPases, and the dynamins, for which several decades of research have indicated robust participation in immune functions [15, 28, 95].

### ParA-like domains in immune functions: unity or diversity?

Research on immune systems across the tree of life has revealed that certain (super)families of domains have a predilection for well-defined roles within these systems (e.g., as sensors, effectors, signal generators or negative regulators). While these are their modal behaviors, there is often greater functional diversity when they are analyzed more closely. Thus, for example, related domains from the TIR/DrHyd/SLOG assemblage of Rossmannoid (especially when encoded by tandem gene arrays) function not only as effectors but also as signal generators and sensors [22, 208, 209]. Likewise, the AGO/PIWI domains act not only as effectors but also as sensors of invasive nucleic acids that regulate additional associated effectors [210–212]. Further, domains from the CARF and SAVED assemblage of Rossmannoid domains function as both sensors and signal-terminating endonucleases, cleaving oligonucleotide messengers [157, 213–215]. Hence, the question arises: what is the primary role of the ParA domains in immune processes, and whether they might have a broader functional horizon, as the above domains?

A key finding of our current study was that the ParA domains have been repeatedly recruited for immune functions within the radiation of the family in core cellular functions (e.g., chromosome segregation; Figure 2A). This points to a certain fungibility in their modes of action between these roles and implies that one could transfer inferences from the well-studied core systems to the defense systems. We posit that this applies to the three basic activities of ParA domains, namely [35, 101, 216, 217]: (i) the ability to act as a switch that utilizes the NTP-hydrolysis cycle; (ii) assembly into multimers; (iii) regulated tethering interactions with partners in macromolecular complexes.

Despite their organizational diversity, we find that a common theme across the ParA-HeH, ParA-wHTH, ParA-CBM, and HinPO conflict systems is a core of two components: the ParA component coupled to a sensor domain (Figures 4B,5C,E,K). Considering this together with the above evolutionary evidence for a unified mode of action, we argue that across these systems, ParA functions not only as an NTP-dependent conformational switch but also as the primary effector, acting via the formation of multimeric assemblies. Such multimeric assemblies could ultimately limit infection by containing the selfish element, i.e., physically blocking the “Central Dogma” processes. Indeed, the conservation of the DNA-binding interface of ParA domains (first identified in the housekeeping ParABS system [81]) across nearly all ParA homologs implicated in immune functions (Supplementary Fig. S5) suggests that direct nucleic acid “coating” by ParA multimers might be central to its action. Importantly, this is also consistent with the growing body of evidence that several immune effectors across the tree of life limit infection by forming large filaments or multimeric complexes, often in a nucleotide/NAD+-dependent manner, upon activation [218–222].

However, keeping with the broadening of the roles we noted for other domains participating in immunity, we observed potential variations on the above primary functional theme of the ParA domains. First, analysis of their architectures revealed that in the TA-ParA and TIR-ParA systems, the ParA domain plays a role comparable to that of STAND NTPases, which have emerged as a major theme in anti-selfish element immunity across the Tree of Life [30]. We had previously shown that in other systems with convergent, broadly “tripartite” architectures, the STAND NTPase might be replaced by an AP-GTPase (a small Ras-like GTPase domain) or an HSP90- or HSP70-like NTPase domain [29, 31]. In all these cases, the distantly related or unrelated NTPase domain is believed to function as a switch that transmits the infection signals recognized by associated sensor domains to the linked apoptotic effector domains. Indeed, in comparable systems with ParA domains, it is well-suited to play a similar role as a switch that activates associated effectors. However, just as the STAND NTPases are known to multimerize [223], ParA could likewise exhibit its basic multimerization activity even in these contexts to form higher order active assemblies. Second, in the ParA-HeH systems, we found that the ParA domains consistently contain a variable insert. In such cases, the variability is a likely marker of an arms race with comparably variable elements on the invader side. Hence, in these cases, the role of the ParA domain could be additionally extended to that of a sensor of invader presence.

### ParA-based immune systems as a surprising marker for bacterial multicellularity

Unexpectedly, several of the ParA systems described here are widespread and significantly enriched among phylogenetically distant prokaryotes with a clear multicellular stage in their life cycle (i.e., over and beyond simple coloniality like biofilms [89, 90]). These primarily include the HinPO, Effector-ParAB, and TIR-ParA systems (Figure 3 and Supplementary Fig. S1), with the last of these closely paralleling the previously reported enrichment of certain STAND NTPase systems in multicellular bacteria [28, 29]. More generally, previous studies have shown that, across the Tree of Life, the emergence of immunity systems that trigger apoptosis went hand-in-hand with multicellularity [28, 29, 34, 224]. We posit that this connection to multicellularity is directly related to the above considerations on ParA functions. First, the multiple ParA systems predicted to initiate apoptosis as their primary mode of action are likely to stabilize incipient multicellularity in a lineage by limiting the spread of an invading selfish element across the cellular ensemble via self-sacrifice. Second, the property of the ParA domains to form multimeric ensembles could be especially apposite in multicellular contexts by forming physical barriers to the transmission of infectious elements to neighboring cells within the ensemble. This is indeed consistent with certain other domains, such as the cytoskeletal tubulin homologs, observed in other immune mechanisms enriched in multicellular bacteria [28].

### ParA domains compared to other P-loop NTPases in immune processes

Given their versatility as NTP-utilizing switches, motors and biosynthetic enzymes, P-loop NTPases are present in practically every cellular process across the Tree of Life, making them one of the largest superfamilies of proteins encoded by the genome of a given cellular organism. The same versatility is also reflected in the P-loop NTPases across systems deployed in biological conflicts, including those involved in immunity [34]. As a result, several major clades of P-loop NTPases have undergone vast radiations associated with several distinct roles in immune mechanisms.

Nevertheless, these can be unified under a few broad themes in each major clade. For example, the helicase clade, with its two major subgroups, the superfamily-1 and superfamily-2 helicases, primarily function as nucleic acid sensors [33] or as motors powering DNA translocation in R-M and allied defense systems [225]. Members of the ABC ATPase clade have also diversified in comparable nucleic-acid-targeting defense systems, but mediate a different set of processes, including sensor roles in detecting nucleotide signals, DNA topology and end sensing [118, 226, 227], conformational switching [228] and long-range DNA translocation [119]. NTPases from the AAA+ clade have colonized a wide range of roles in immunity. Some, like those of the McrB subclade, function comparably to helicases in DNA translocation [229]. Others, like those from the MoxR subclade, act as chaperones that facilitate the assembly of functional immunity-related complexes[28]. Yet others, namely the STAND NTPases from the Orc-Cdc6 subclade, act as conformational switches that activate associated effector domains upon sensing of viral infection [30, 230]. TRAFAC GTPases participating in immune roles fundamentally act as switches, utilizing this to function either as effectors themselves or as regulators of effector activation comparable to the STAND branch of AAA+ NTPases [15, 28, 95, 231, 232].

Here, we present evidence that the ParA clade of NTPases from the SIMIBI subgroup of the GTPase clade has evolved functions that are broadly comparable to those of other P-loop NTPases. While they appear to use ATP as their primary substrate by virtue of the degeneration of the [TN]KxD motif associated with GTP recognition, they are still predicted to function as switches comparable to the versions in institutionalized cellular roles (Figure 5B, F-G) [101]. In this capacity, they are predicted to act comparably to the STAND NTPases and TRAFAC GTPases in immune mechanisms. Given that the deployment of potentially fatal effectors is a life-or-death cellular decision, their regulation has provided a fertile functional niche for distantly related P-loop NTPases lines with a comparable capacity for NTP-dependent switching. We also propose that the ParA domains play a distinct role in the assembly of multimeric effector complexes, in partnership with an associated invader-sensing component, and in direct interactions with DNA. This appears to be a key distinguishing feature of the recruitment of ParA domains compared with most other P-loop NTPases recruited to immune functions.

### Conclusions

Using sensitive sequence and structure analysis, combined with contextual information, we identify ParA domains as a ‘fulcrum’ domain in a wealth of previously unexplored or poorly studied immune systems. While these are primarily found in prokaryotes, we also find evidence of their transfer to photosynthetic eukaryotes and certain metazoans on multiple occasions. This study establishes the ParA domain as a distinct paradigm among the P-loop NTPases deployed in immune mechanisms. While it shares functional features with TRAFAC GTPases and STAND NTPases, we also establish its distinct effector role, potentially involving partnership with a wide range of sensor and secondary effector domains. The functional partners associated with ParA in these systems also served as anchors for uncovering new potential immune systems utilizing a wide cast of underappreciated protein domains and potential novel mechanisms. We hope the conceptual frameworks presented in this work will help guide future wet-lab studies of immune processes and open the door to the development of novel reagents.

## Data availability

All data supporting the findings of this study are available within the article and its Supplementary Data.

## Supplementary data

Supplementary Data are available at NAR Online

## Author contributions

**Huan Li:** Conceptualization; data curation; formal analysis; investigation; methodology; visualization; writing–original draft. **Maxwell Burroughs:** Data curation; formal analysis; investigation; methodology; validation; writing–original draft. **L Aravind:** Conceptualization; formal analysis; funding acquisition; investigation; methodology; project administration; supervision; validation; writing–review and editing.

