## Supplementary_Figures for "Widespread Recruitment of ParA-like NTPases into Prokaryotic Immunity reveals a diversity of anti-selfish element strategies"

This file contains: Supplementary figures S1-S9

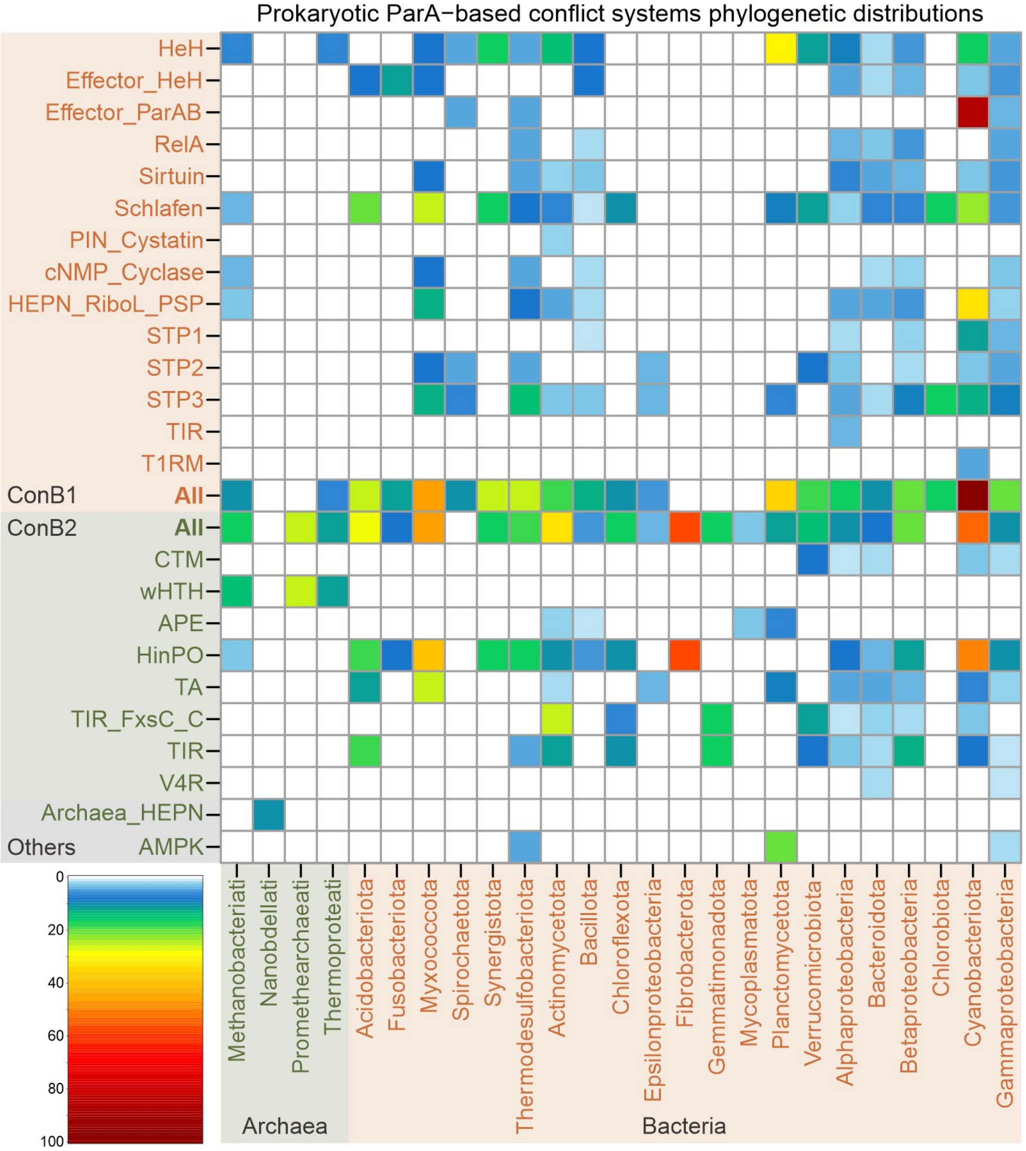


**Figure S1:** Phyletic patterns of prokaryotic ParA-based biological conflict systems. The x-axis shows taxon names, grouped into the Archaea and Bacteria superkingdoms. The y-axis lists system names, including those in ConB1, ConB2, and other groups. Grid colors indicate the percentage of representation within each specified taxon.

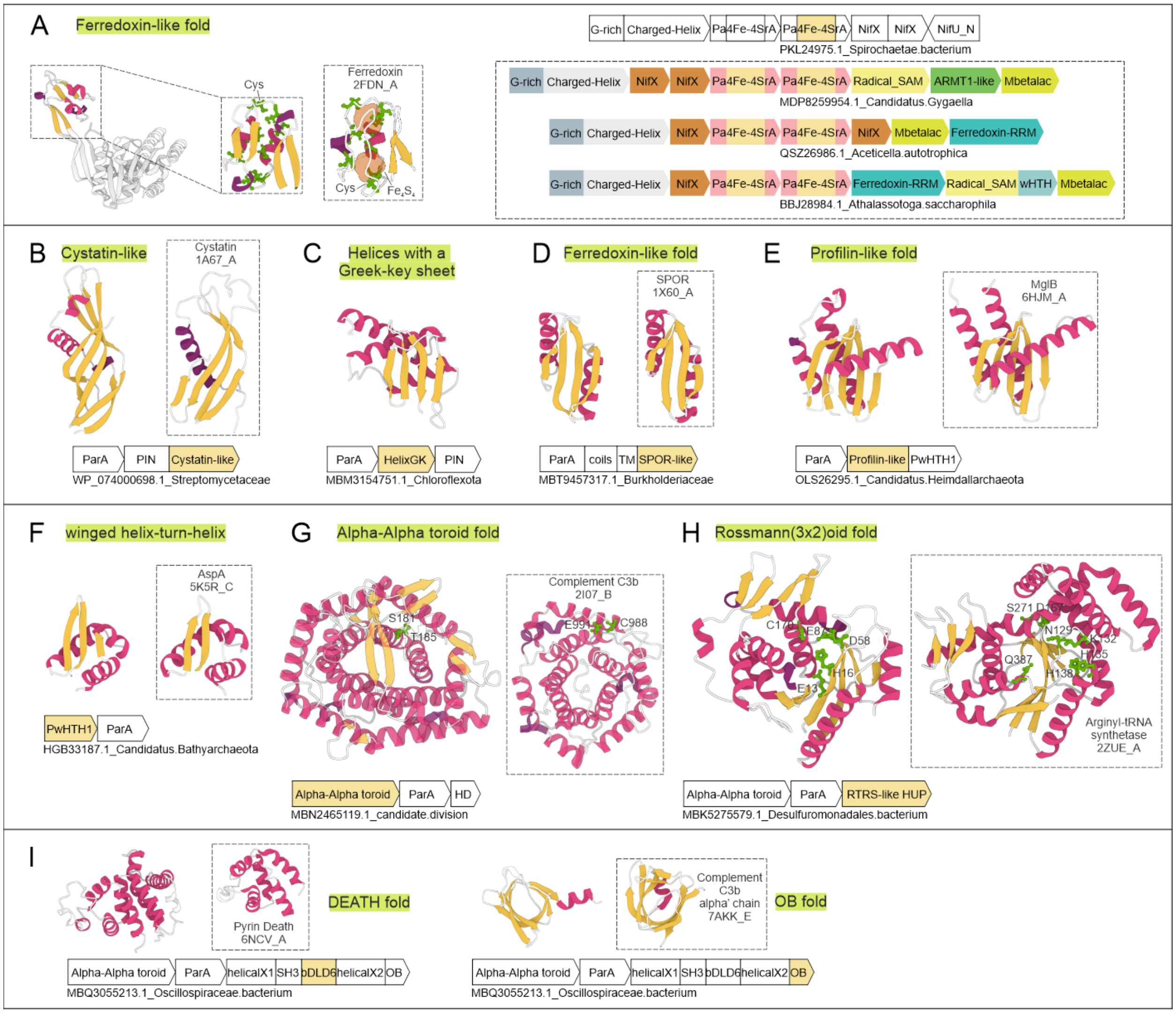


**Figure S2**: Cartoon representations of structural models for novel domains identified in this study. (A) The 4Fe-4S domain. Representative variants of ParA homologs involved in nitrogen fixation pathways with additional domains are also shown here. (B) The Cystatin-like domain. (C) The HelixGK domain. (D) The SPOR-like domain. (E) The Profilin-like domain. (F) The PwHTH1 domain. (G) The Alpha-Alpha toroid domain. (H) The RTRS-like domain. (I) The bDLD6 domain and the OB-like domain. A depiction of the corresponding system for each domain is shown below, including the NCBI ID and the organism name. A distant structural homolog, with its name and PDB ID, for each domain is indicated within a dashed box. For enzymatic domains, conserved residues predicted for catalytic activity are shown as ball-and-stick models with labels. The fold shared by the domain and its distant homolog is also shown above or alongside.

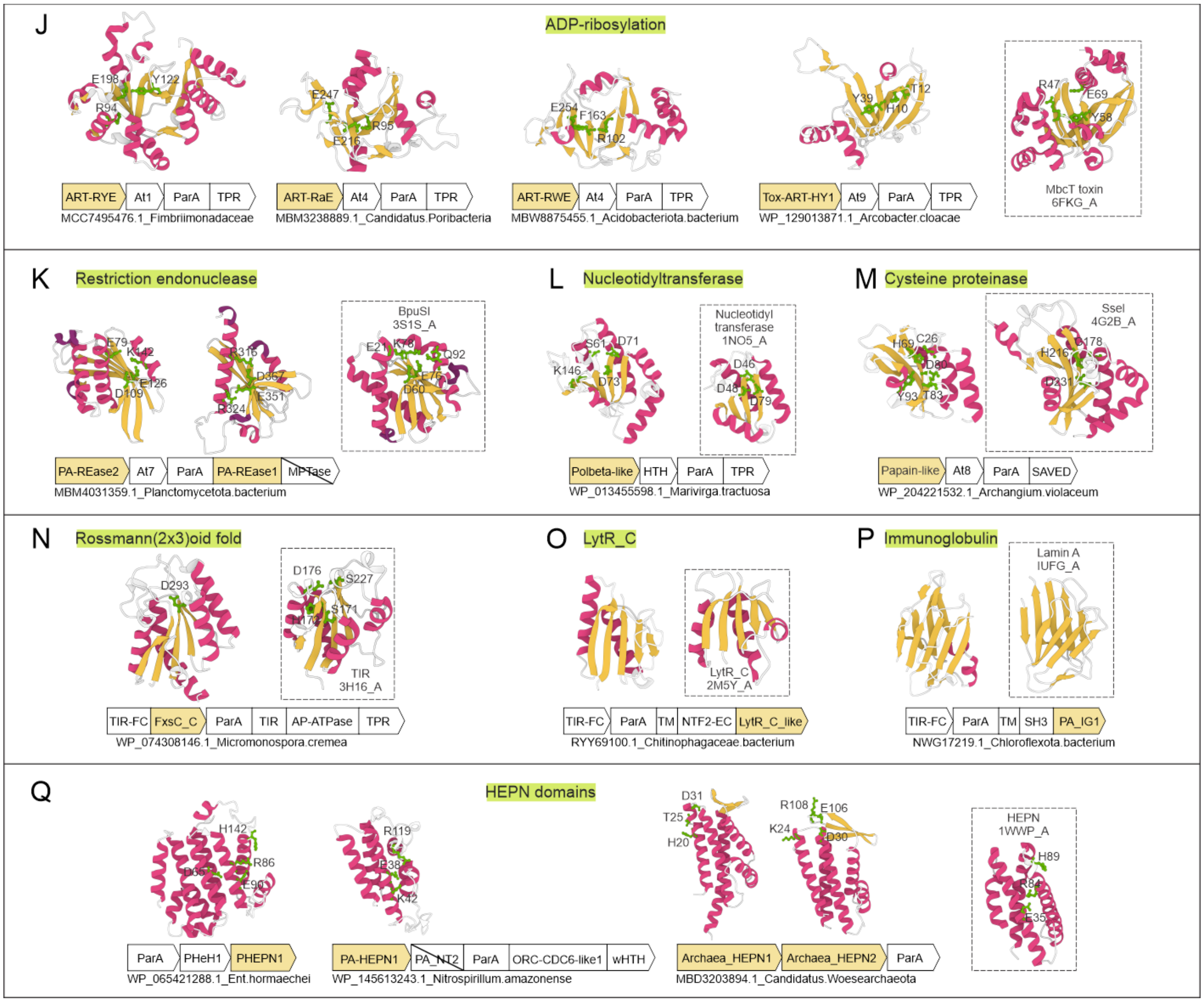


**Figure S2 (continued)**: Cartoon representations of structural models for novel domains identified in this study. (J) The novel ART domains. (K) The novel restriction endonuclease domains. (L) The Polbeta-like domain. (M) The Papain-like domain. (N) The FxsC_C domain. (O) The LytR_C-like domain. (P) The PA_IG1 domain. (Q) The novel HEPN domains.

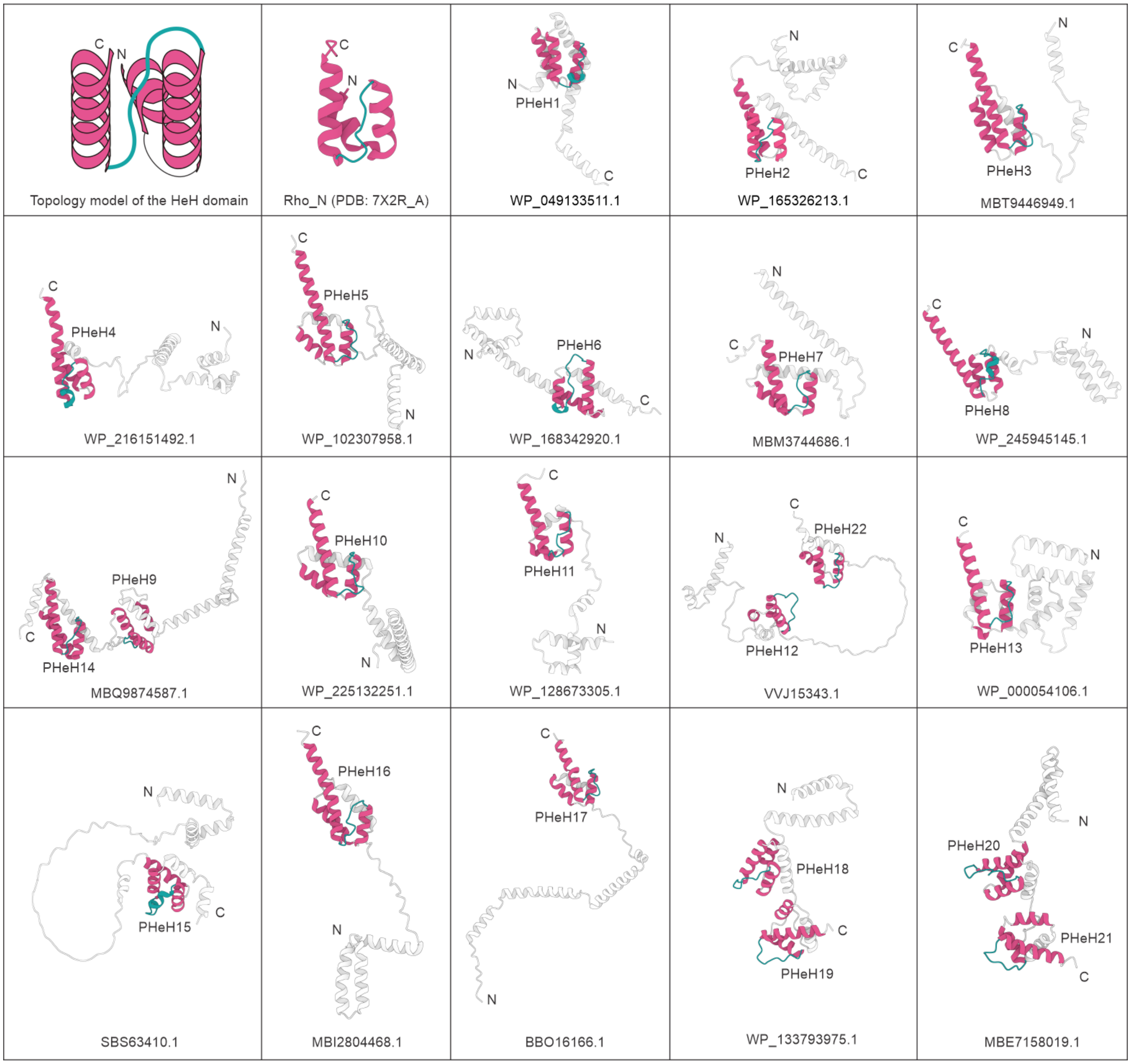


**Figure S3**: Diverse PHeH domains in the ParA-HeH system. The first grid shows a topology model of the HeH-type domain, and the second grid shows the structure of a classical HeH domain. The remaining grids show cartoon representations of proteins containing PHeH domains in the ParA-HeH system. Only HeH domains are colored.

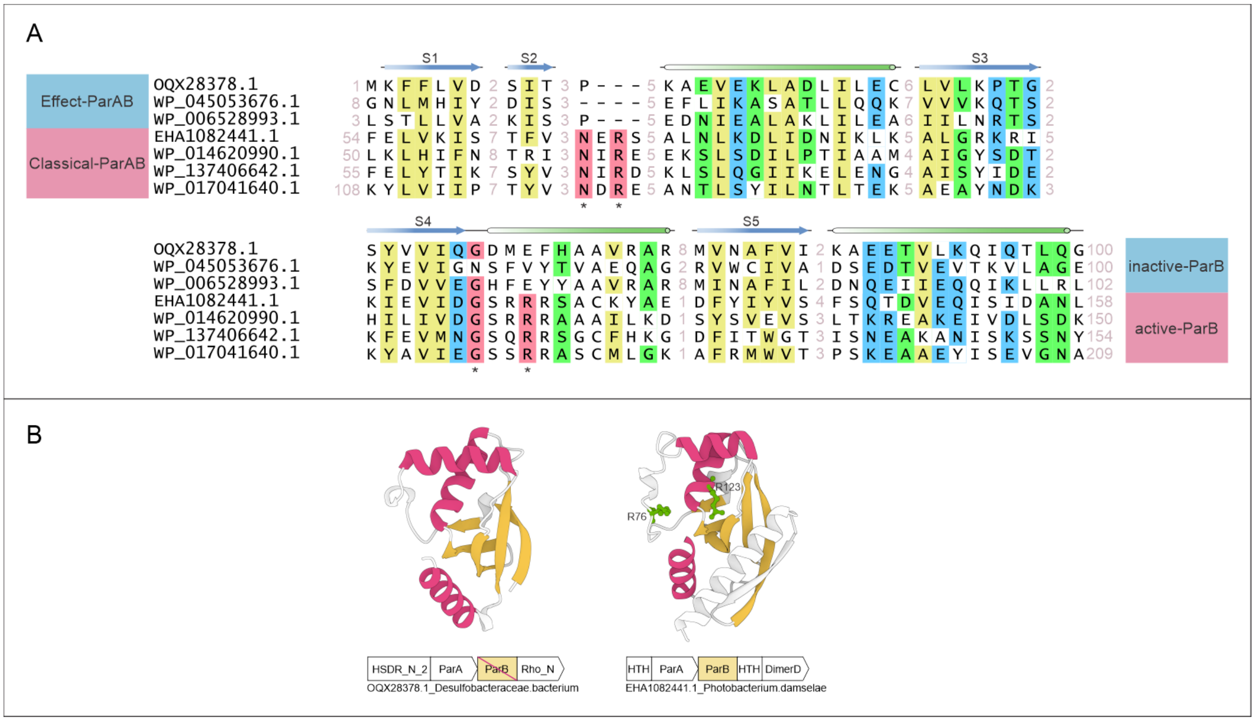


**Figure S4**: ParB domains in the effector-ParAB system are inactive. (A) Multiple sequence alignment of ParB domains from the effector-ParAB system (upper three) and the classical ParABS system (bottom four). Conserved hydrophobic columns are colored yellow; conserved polar columns are colored blue; conserved small columns are colored green; and conserved residues in ParB domains from the classical ParABS system are colored red and indicated by asterisks. Schematic representations of secondary structures are shown above. (B) Cartoon representations of structural models for a ParB domain from the effector-ParAB system and a ParB domain from the classical ParABS system. Conserved catalytic residues in the latter are shown as ball-and-stick models. A depiction of the corresponding system for each ParB is shown below, including the NCBI ID and the organism name.

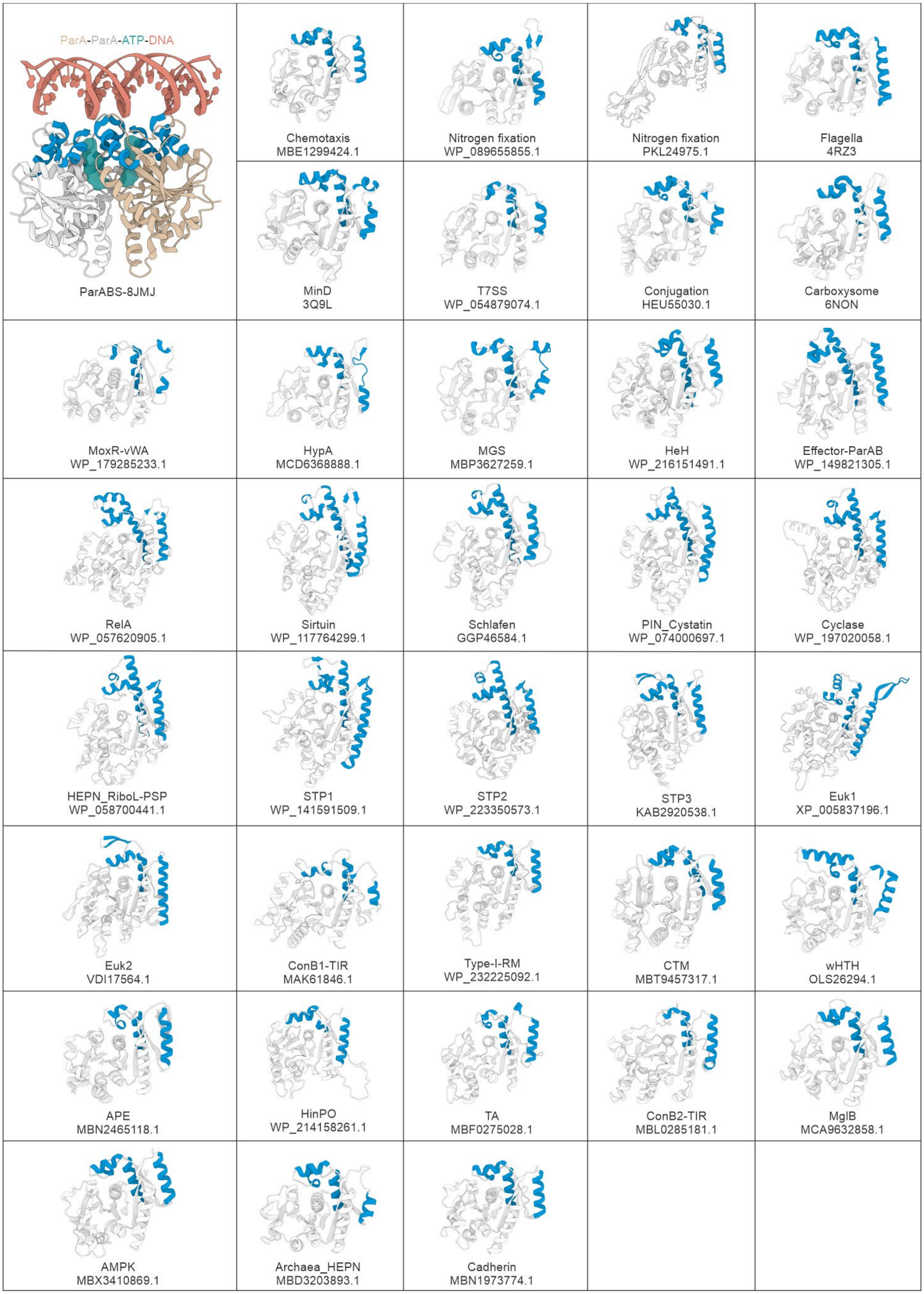
ParA-ParA-ATP-DNA
ParABS-8JMJ
Chemotaxis
MBE1299424.1
Nitrogen fixation
WP_089655855.1
Nitrogen fixation
PKL24975.1
Flagella
4RZ3
MinD
3Q9L
T7SS
WP_054879074.1
Conjugation
HEU55030.1
Carboxysome
6NON
MoxR-vWA
WP_179285233.1
HypA
MCD6368888.1
MGS
MBP3627259.1
HeH
WP_216151491.1
Effector-ParAB
WP_149821305.1
RelA
WP_057620905.1
Sirtuin
WP_117764299.1
Schlafen
GGP46584.1
PIN_Cystatin
WP_074000697.1
Cyclase
WP_197020058.1
HEPN_RiboL-PSP
WP_058700441.1
STP1
WP_141591509.1
STP2
WP_223350573.1
STP3
KAB2920538.1
Euk1
XP_005837196.1
Euk2
VDI17564.1
ConB1-TIR
MAK61846.1
Type-I-RM
WP_232225092.1
CTM
MBT9457317.1
wHTH
OLS26294.1
APE
MBN2465118.1
HinPO
WP_214158261.1
TA
MBF0275028.1
ConB2-TIR
MBL0285181.1
MglB
MCA9632858.1
AMPK
MBX3410869.1
Archaea_HEPN
MBD3203893.1
Cadherin
MBN1973774.1

**Figure S5**: DNA-binding elements are conserved across the ParA family. The first grid depicts a structural model of the ATP-ParA-DNA complex, in which the ParA protein originates from a classical ParABS system. The remaining grids show cartoon representations of ParA domains from different systems, including fundamental cellular processes and biological conflict systems. Counterparts of DNA-binding elements are colored blue in all models. NCBI or PDB IDs are shown below.

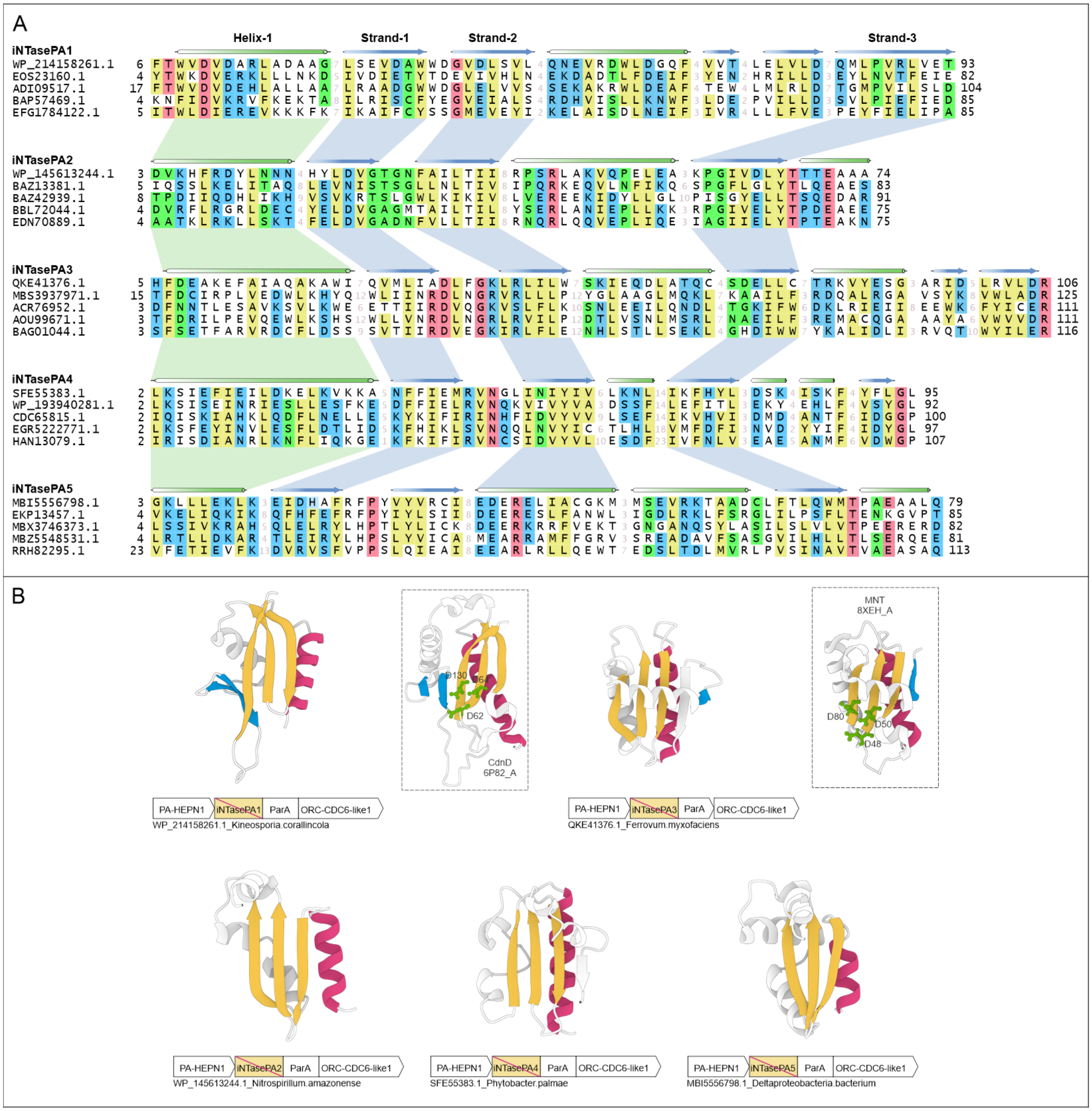


**Figure S6**: iNTasePA1 to iNTasePA5 from the HinPO systems are inactive nucleotidyltransferase domains. (A) Multiple sequence alignment of the iNTasePA1 to iNTasePA5 domains. Corresponding core elements across different domains are connected. (B) Cartoon representations of structural models for iNTasePA1 to iNTasePA5 domains. Only core elements are colored.

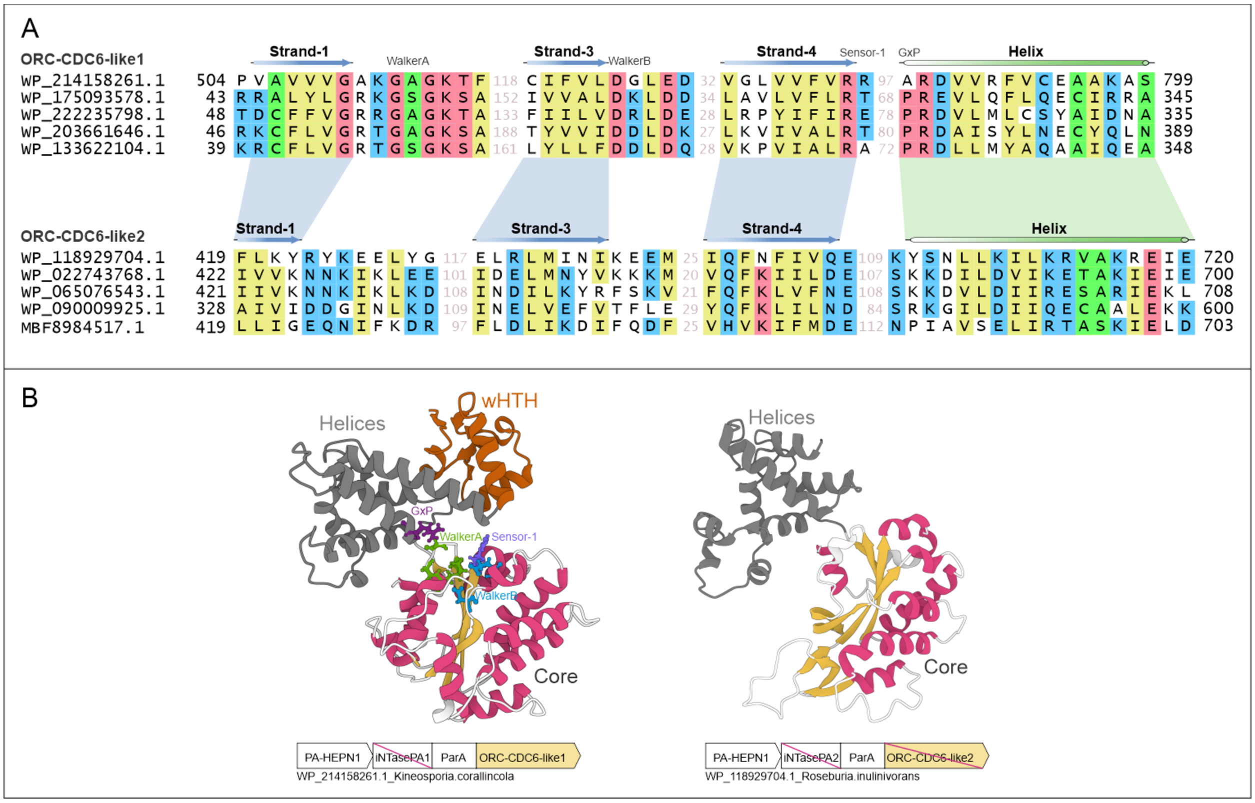


**Figure S7**: ORC-CDC6-like1 (active) and ORC-CDC6-like2 (inactive) are STAND NTPases in the HinPO system. (A) Multiple sequence alignment of the ORC-CDC6-like1and ORC-CDC6-like2 domains. Corresponding core elements across different domains are connected. (B) Cartoon representations of structural models for ORC-CDC6-like1 and ORC-CDC6-like2 domains.

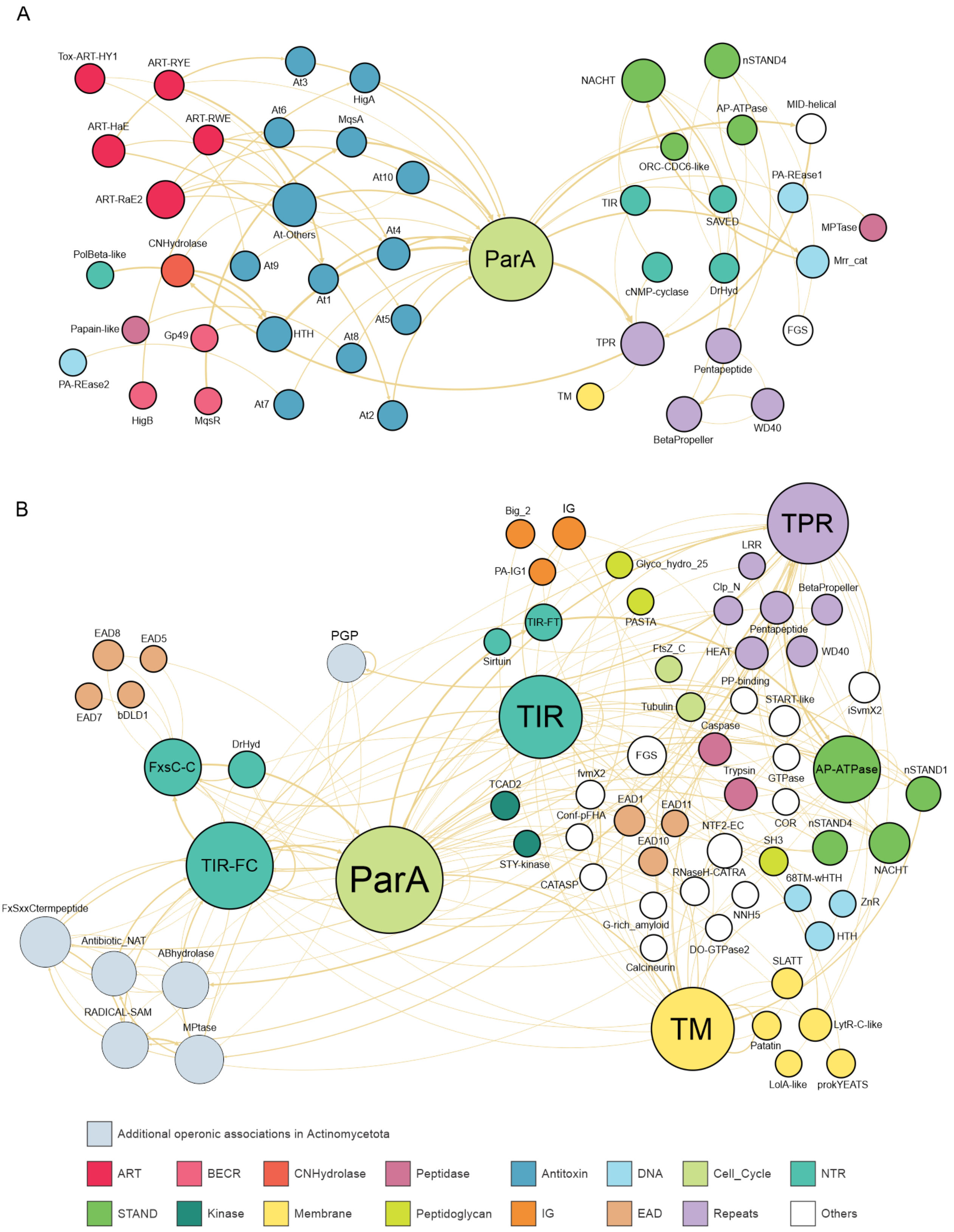
A
Tox-ART-HY1
ART-RYE
At3
HigA
ART-HaE
ART-RWE
At6
MqsA
At10
ART-RaE2
At-Others
ParA
PolBeta-like
CNHydrolase
At9
At4
At1
Papain-like
Gp49
HTH
At8
At5
PA-REase2
HigB
MqsR
At7
At2
NACHT
nSTAND4
AP-ATPase
MID-helical
ORC-CDC6-like
PA-REase1
TIR
SAVED
MPTase
cNMP-cyclase
DrHyd
Mrr_cat
FGS
TPR
Pentapeptide
TM
BetaPropeller
WD40
B
Big_2
IG
PA-IG1
Glyco_hydro_25
TPR
LRR
Clp_N
BetaPropeller
PASTA
TIR-FT
Sirtuin
Pentapeptide
EAD8
EAD5
PGP
FtsZ_C
HEAT
WD40
EAD7
bDLD1
PP-binding
iSvmX2
START-like
Tubulin
TIR
Caspase
DrHyd
FxsC-C
FGS
Trypsin
GTPase
AP-ATPase
nSTAND1
TCAD2
fvmX2
EAD1
EAD11
COR
nSTAND4
Conf-pFHA
NTF2-EC
SH3
NACHT
TIR-FC
ParA
STY-kinase
EAD10
CATASP
RNaseH-CATRA
68TM-wHTH
ZnR
FxSxxCtermpeptide
G-rich_amyloid
NNH5
Antibiotic_NAT
ABhydrolase
DO-GTPase2
HTH
Calcineurin
SLATT
RADICAL-SAM
MPtase
TM
Patatin
LytR-C-like
LolA-like
prokYEATS
Additional operonic associations in Actinomycetota
ART
BECR
CNHydrolase
Peptidase
Antitoxin
DNA
Cell_Cycle
NTR
STAND
Kinase
Membrane
Peptidoglycan
IG
EAD
Repeats
Others

**Figure S8**: Domain networks of apoptosis-like systems, including (A) TA-related systems and (B) TIR-related systems. Node size is scaled by degree, and edge thickness is scaled by occurrence frequency. Node colors represent domain functional groups, as indicated below.

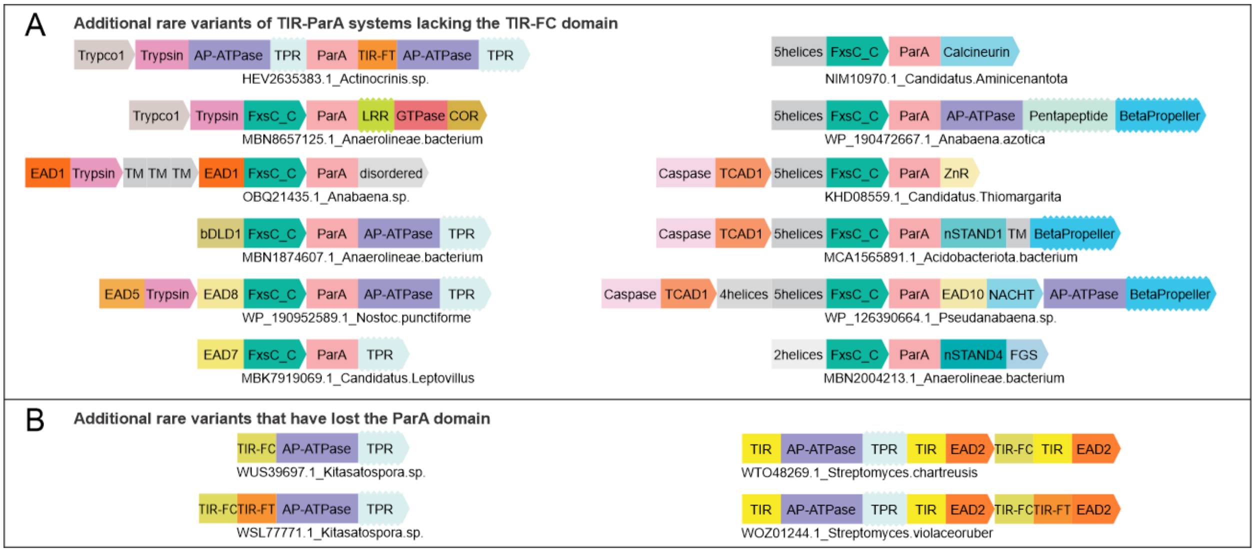


**Figure S9**: (A) Additional rare variants of TIR-ParA systems lacking the TIR-FC domain, and (B) Additional rare variants that have lost the ParA domain.
